# Single-Cell Transcriptomic Profiling of the Zebrafish Inner Ear Reveals Molecularly Distinct Hair Cell and Supporting Cell Subtypes

**DOI:** 10.1101/2022.09.08.507120

**Authors:** Tuo Shi, Marielle O. Beaulieu, Lauren M. Saunders, Peter Fabian, Cole Trapnell, Neil Segil, J. Gage Crump, David W. Raible

## Abstract

A major cause of human deafness and vestibular dysfunction is permanent loss of the mechanosensory hair cells of the inner ear. In non-mammalian vertebrates such as zebrafish, regeneration of missing hair cells can occur throughout life. While a comparative approach has the potential to reveal the basis of such differential regenerative ability, the degree to which the inner ears of fish and mammals share common hair cells and supporting cell types remains unresolved. Here we perform single-cell RNA sequencing of the zebrafish inner ear at embryonic through adult stages to catalog the diversity of hair cell and non-sensory supporting cells. We identify a putative progenitor population for hair cells and supporting cells, as well as distinct hair cells and supporting cell types in the maculae versus cristae. The hair cell and supporting cell types differ from those described for the lateral line system, a distributed mechanosensory organ in zebrafish in which most studies of hair cell regeneration have been conducted. In the maculae, we identify two subtypes of hair cells that share gene expression with mammalian striolar or extrastriolar hair cells. In situ hybridization reveals that these hair cell subtypes occupy distinct spatial domains within the two major macular organs, the utricle and saccule, consistent with the reported distinct electrophysiological properties of hair cells within these domains. These findings suggest that primitive specialization of spatially distinct striolar and extrastriolar hair cells likely arose in the last common ancestor of fish and mammals. The similarities of inner ear cell type composition between fish and mammals also support using zebrafish as a relevant model for understanding inner ear-specific hair cell function and regeneration.

## Introduction

Mechanosensory hair cells of the inner ear are responsible for sensing sound and head position in vertebrates. Hair cells are notoriously susceptible to damage from multiple types of insults, including noise and ototoxic drug exposure. Studies of hair cell physiology in mammals are limited by the location of the inner ear within the temporal bone, which precludes many targeted manipulations and in vivo imaging beyond the neonatal stage. As a result, non-mammalian vertebrates with analogous, more easily accessible hair cells have become useful models for studying hair cell development, death, and regeneration. Non-mammalian vertebrates such as birds and fish can regenerate hair cells of the auditory and vestibular systems that are lost due to injury (Stone and Cotanche, 2007; Monroe et al., 2015). This differs from mammals, where cochlear hair cell death leads to permanent hearing loss (Corwin and Cotanche, 1988; Yamasoba and Kondo, 2006), and limited regeneration of vestibular hair cells results in minimal recovery of function (Golub et al., 2012). Non-mammalian model systems of hair cell regeneration have the potential to reveal conserved pathways that can be targeted to promote hair cell survival and regeneration in humans. However, the extent of hair cell molecular homology across vertebrates remains unclear.

Due to its accessibility for manipulation and imaging, the zebrafish lateral line system has been widely used to study mechanisms of hair cell physiology (Pickett and Raible, 2019; Sheets et al., 2021). The lateral line is an external sensory system that allows aquatic vertebrates to detect local movement of water. Sensory organs of the lateral line, called neuromasts, contain hair cells and supporting cells that share properties with those of the inner ear. However, relative to the lateral line, cells in the zebrafish inner ear are likely more similar to their mammalian counterparts, raising the potential for it to be a more comparable system in which to study hair cell function.

Zebrafish and mammals share several inner ear sensory organs. Three semicircular canals with sensory end organs called cristae sense angular rotation of the head. Two additional sensory end organs detect linear acceleration and gravity: the utricular and saccular macula with associated otolith crystals (Figure 1). Fish lack a specific auditory structure such as the mammalian cochlea and instead sense sound through the saccule, utricle, and a third otolith organ, the lagena.

**Figure 1.**
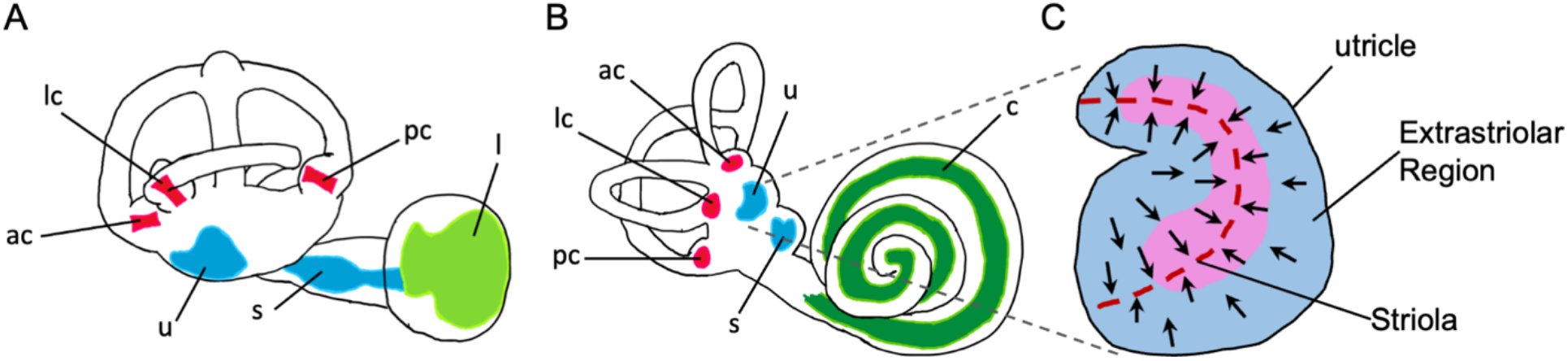
Anatomy of zebrafish and mouse inner ears. Illustrations of zebrafish A) and mouse B) inner ears showing homologous end organs in the semicircular canal crista ampullaris (red) and macula otolith organs (blue). Light green and dark green represent unique end organs of the lagena in the zebrafish and cochlea in mice. C) Illustration of the mouse utricle showing striolar and extrastriolar regions of the sensory organ. Arrows represent hair cell planar polarity within the sensory organ and red dashed line represents the line of polarity reversal within the striola. ac: anterior crista, c: cochlea, l: lagena, lc: lateral crista, pc: posterior crista, s: saccule, u: utricle.

Although historically the saccule and utricle were thought to be for vestibular function and the lagena analogous to the cochlea for sound detection, there is now substantial evidence for all three otolith end organs being used for sound detection with diverse specializations across fishes (Popper and Fay, 1993). In larval zebrafish, both saccule and utricle hair cells respond to sound stimuli of frequencies between 100-4000 Hz (Yao et al., 2016; Privat et al., 2019; Favre-Bulle et al., 2020).

Within the mammalian utricle and saccule, there are both morphological and spatial differences between hair cells (Lysakowski and Goldberg, 2004; Eatock and Songer, 2011). Hair cells are broadly classified by their morphology and innervation, with Type I hair cells having calyx synapses surrounding the hair cell body and Type II hair cells having bouton synapses. Both Type I and Type II cells can be found within the central region of the macular organs known as the striola and in the surrounding extrastriolar zones. Although the role of spatial segregation into striolar versus extrastriolar zones has not been fully elucidated, across these regions hair cells vary in morphology, electrophysiology, and synaptic structure(Desai et al., 2005; Li et al., 2008). The striola is characterized by hair cells with taller ciliary bundles and encompasses a line of polarity reversal where hair cells change their stereocilia orientation (Figure 1C). Whereas distinct Type I and Type II hair cells, and in particular the calyx synapses typical of Type I cells, have not been identified in fishes, spatial heterogeneity in the maculae, including those of zebrafish, has been previously noted (Chang et al., 1992; Platt, 1993; Popper, 2000; Liu et al., 2022). However the homologies of cells at the cellular and molecular level have remained unknown.

Recent single-cell and single-nucleus RNA-sequencing efforts have generated a wealth of transcriptomic data from hair cells in several model systems, facilitating more direct comparison of cell types and gene regulatory networks between species. Although single-cell transcriptomic data have recently been published for the zebrafish inner ear (Jimenez et al., 2022; Qian et al., 2022), the diversity of hair cell and supporting cell subtypes has not been thoroughly analyzed. In order to better understand the diversification of cell types in the zebrafish inner ear, and their relationships to those in mammals, here we perform single-cell RNA sequencing (scRNA-seq) of the zebrafish inner ear from embryonic through adult stages. We find that hair cells and supporting cells from the zebrafish inner ear and lateral line are transcriptionally distinct, and that hair cells and supporting cells differ between the cristae and maculae. All of these distinct cell types are present during larval development and are maintained into adulthood. In situ hybridization reveals that these hair cell subtypes occupy distinct spatial domains within the utricle and saccule, and computational comparison of hair cell types reveals homology with striolar and extrastriolar hair cell types in mammals. These findings point to an origin of striolar and extrastriolar hair cell types in at least the last common ancestor of fish and mammals.

## Results

### Inner ear hair cells and supporting cells are distinct from those of the lateral line

To assess differences between inner ear and lateral line cells, we analyzed a subset of cells from a large single-nucleus RNA-seq dataset of whole zebrafish at embryonic and larval stages (24-96 hours post-fertilization (hpf)), which was prepared by single nucleus combinatorial indexing and sequencing (“sci-Seq”; Saunders et al., 2022). A total of 16,517 inner ear and lateral line cells were combined and re-processed using Monocle 3 (Figure 2A-B). Inner ear nonsensory cells were identified by expression of known marker genes, including the transcription factor gene *sox10* (Dutton et al., 2009) in combination with inner ear supporting cell genes *(stm, otog, otogl, otomp, tecta* and *oc90*; Figure 2C) (Söllner et al., 2003; Kalka et al., 2019; Petko et al., 2008; Stooke-Vaughan et al., 2015). Lateral line nonsensory cells were identified by expression of known markers *fat1b, tfap2a, tnfsf10l3, lef1, cxcr4b, fgfr1a* and *hmx3a* (Figure 2D) (Steiner et al., 2014; Thomas and Raible, 2019; McGraw et al., 2011; Haas and Gilmour, 2006; Lee et al., 2016; Feng and Xu, 2010). We identified hair cells by expression of the pan-hair cell genes *otofb, cdh23, pcdh15a, ush1c, myo7aa, slc17a8 and cacna1da* (Figure 2E) (Chatterjee et al., 2015; Söllner et al., 2004; Seiler et al., 2005; Phillips et al., 2011; Ernest et al., 2000; Obholzer et al., 2008; Sheets et al., 2012). To distinguish between inner ear and lateral line hair cells, we queried expression of previously described markers for inner ear (*gpx2, kifl, strc*, and *lhfpl5a*) and lateral line (*strc1, lhfpl5b*, and *s100t*) (Erickson et al., 2019; Erickson and Nicolson, 2015). Although many of these markers are at low abundance, these populations are marked distinctly by *strc* and *s100t* (Figure 2E,F).

**Figure 2.**
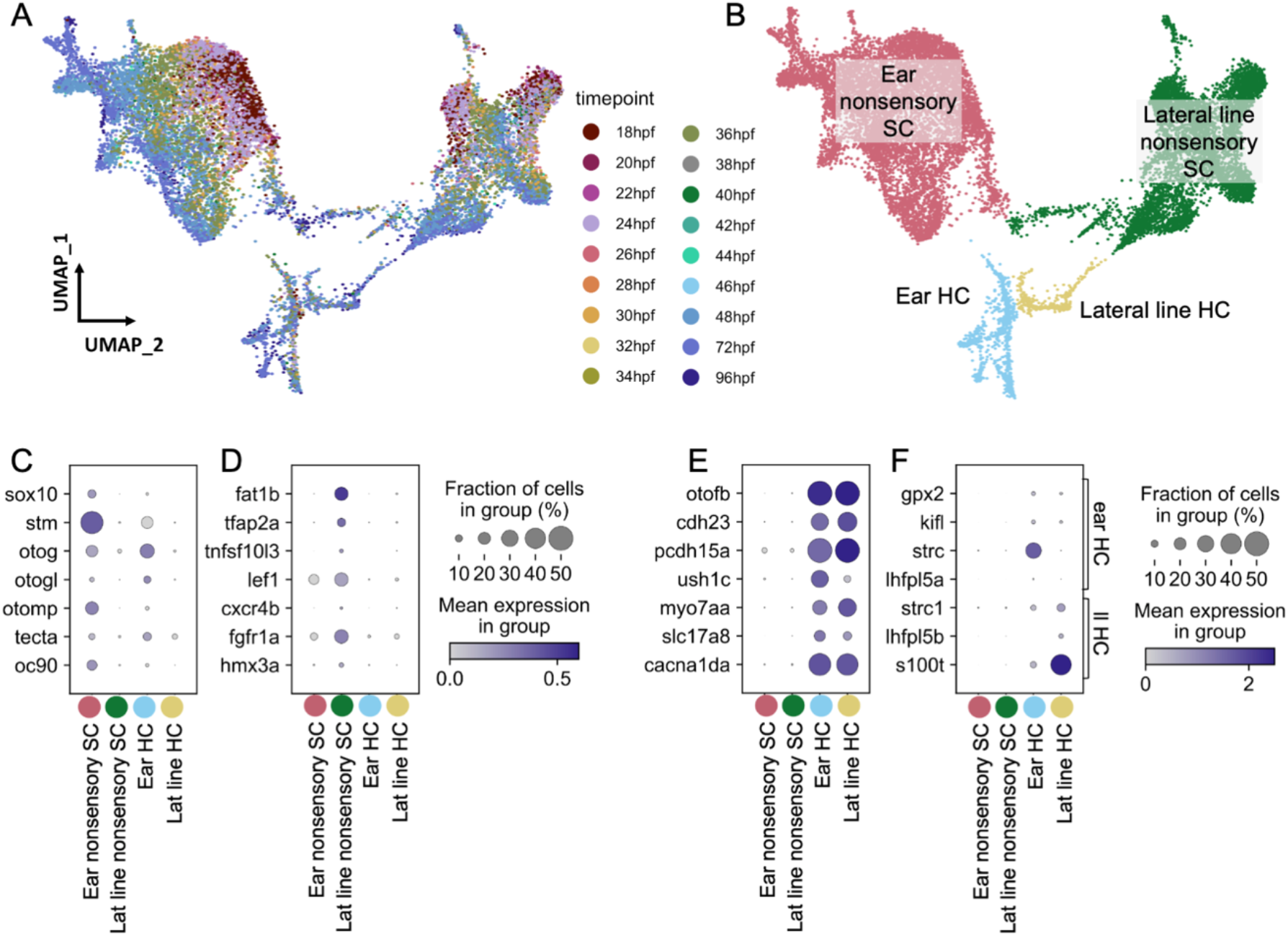
Zebrafish inner ear hair cells and supporting cells are molecularly distinct from those found in the lateral line. Ear and lateral line cells were selected from a whole-embryo single-nucleus RNA-seq dataset from animals between 18 and 96 hpf using known marker genes for hair cells and supporting cells. A-B) snRNA-seq UMAP projection of inner ear and lateral line cells grouped by A) developmental timepoint and B) broad cell type. Widely accepted marker genes for C) inner ear nonsensory cells, D) lateral line nonsensory cells, and E) hair cells show enriched expression in the corresponding clusters from B, confirming their identity. F) Expression of previously identified marker genes for inner ear or lateral line hair cells was used to identify hair cell origin.

Both hair cells and nonsensory supporting cells from the inner ear and lateral line formed distinct clusters, with nonsensory cells from the two mechanosensory organs showing greater distinction than hair cells (Figure 2B, Supplemental Figure 1A). To confirm the relative differences between inner ear and lateral line hair cells and non-sensory cells, PAGA analysis was used to measure the connectivity of clusters (Wolf et al., 2019). This analysis revealed strong connectivity within inner ear supporting cell clusters and within lateral line supporting cell clusters but little connectivity between them (Supplemental Figure 1A, Supplemental Table 1).

The inner ear nonsensory cluster includes structural cells forming the otic capsule, identified by expression of the collagen *col2a1a* and *matrilin4* (*matn4*), indicative of the collagen-containing extracellular matrix (Xu et al., 2018), as well as sensory supporting cells expressing *lfng* (Figure 3D; Supplemental Figure 1B). Inner ear and lateral line supporting cells remain as distinct clusters even when structural *matn4+* cells are excluded from analysis (Supplemental Figure 1C). Thus, both hair cells and supporting cells have distinct gene expression profiles between the inner ear and lateral line at embryonic and larval stages.

**Figure 3.**
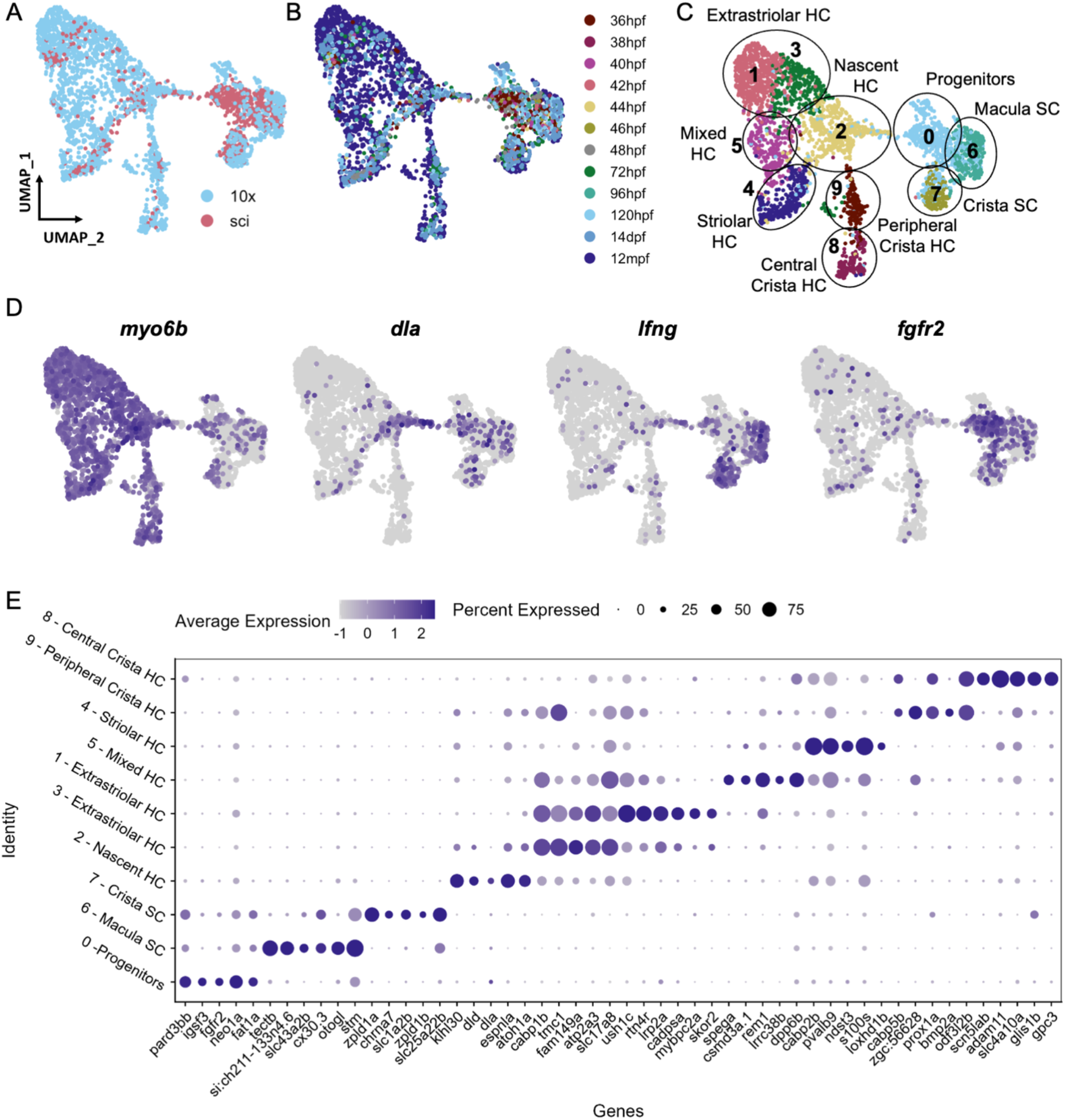
Cell subtypes in the zebrafish inner ear end organs. A-D) Integration and analysis of single cell RNAseq data generated by sci-Seq (sci) or 10x Chromium sequencing (10x) from inner ear hair cells and supporting cells from embryonic (sci), larval (sci,10x) and adult (10x) stages. UMAP projection of cells are grouped by A) dataset of origin and B) timepoint. C) Unsupervised clustering divides cells into 10 clusters that were grouped into 9 cell subtypes. D) Feature plots showing hair cell marker *myo6b*, nascent hair cell marker *dla*, supporting cell marker *lfng* and putative progenitor marker *fgf2b* expression in the integrated dataset. E) Differentially expressed genes across the 10 cell clusters.

### scRNA-seq reveals distinct hair cell and supporting cell populations in the juvenile and adult inner ear of zebrafish

To identify distinct subtypes of inner ear hair cells and supporting cells through adult stages, we performed additional RNA sequencing of single cells isolated from larval (72 and 120 hpf) (Fabian et al., 2022), juvenile (14 days post-fertilization (dpf)), and adult (12 months post-fertilization (mpf)) inner ears. We labeled otic placode cells and their descendants with *Sox10*:Cre to induce recombination of an ubiquitous *ubb*:LOXP-EGFP-LOXP-mCherry transgene (Kague et al., 2012), and then dissected and dissociated ears (14 dpf and 12 mpf) for fluorescence-activated cell sorting (FACS) to purify mCherry+ cells. After construction of scRNA-seq libraries using 10x Chromium technology, hair cells and supporting cells were identified for further analysis based on the expression of hair cell markers *myo6b* and *strc* and supporting cell markers *stm* and *lfng*; structural cells were removed from further analysis based on expression of *matn4* and *col2a1a* (Supplemental Figure 2). Using Seurat, we integrated this dataset with the sci-Seq embryonic and larval dataset (36-96 hpf) (Figure 3A, B). The combined dataset comprises 3246 inner ear cells separated into 10 groups based on unsupervised clustering, with differentially expressed genes for each cluster shown in Figure 3E and Supplemental Table 2. We identified 6 clusters of hair cells based on shared expression of *myo6b, strc*, and *lhfpl5a*, and *gfi1aa* (Yu et al., 2020), a nascent hair cell cluster based on expression of *atoh1a* (Millimaki et al., 2007) and the Notch ligand *dla* (Riley et al., 1999), and two clusters of supporting cells based on expression of *lfng* and *stm* (Figure 3C, D, Supplemental Figure 3). An additional putative progenitor cluster, enriched for cells from embryonic stages, shows weak expression for both hair cell and supporting cell markers, as well as novel genes such as *fgfr2* (Rohs et al., 2013), *fat1a* (Down et al, 2005), *igsf3* and *pard3bb*.

### Putative bipotent progenitor population in the developing inner ear

To understand potential lineage relationships of these clusters, we next performed pseudotime trajectory analysis using Monocle3. We anchored the pseudotime projection at the putative progenitor cell cluster. Analysis revealed two major trajectories toward hair cells and supporting cell clusters (Figure 4A), with distinct patterns of gene expression along each trajectory (Supplemental Table 3). We find that gene expression of the putative progenitor markers follow two patterns: decreasing along both hair cell and supporting cell trajectories (*fgfr2* and *igsf3*) and decreasing only along the hair cell trajectory (*fat1a* and *pard3bb*) (Figure 4C,D). The hair cell trajectory progresses first through the nascent hair cell cluster enriched for early hair cell markers *atoh1a* and *dla* (Cluster 2, Figure 3D, Supplemental Figure 3D), and we observe transient expression of these genes (Figure 4E), as along with increasing expression of hair cell genes *gfi1aa* and *myo6b* (Figure 4F). Along the supporting cell trajectory we observed upregulation of supporting cell-specific markers, including *stm* and *lfng* (Figure 4G). These results support the idea that a population of bipotent progenitors give rise to both hair cells and supporting cells, although future in vivo lineage tracing will be required to test this definitively.

**Figure 4.**
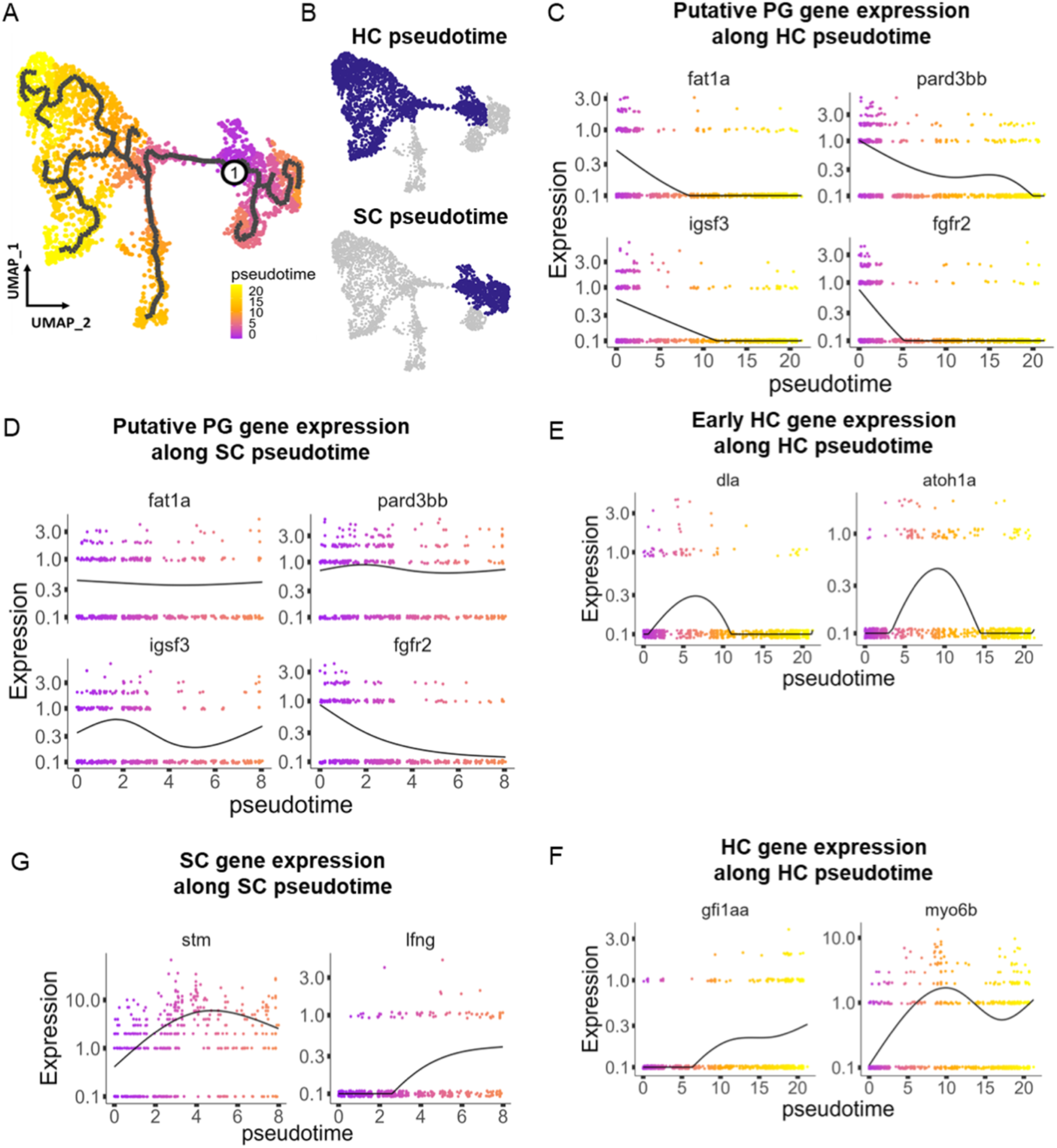
Pseudotime analysis reveals potential hair cells and supporting cell bipotent progenitor population in the zebrafish inner ear. A) Pseudotime analysis showing simulated developmental trajectories of a putative bipotent progenitor population into both B) macular hair cell and supporting cell clusters. C,D) Changes in putative progenitor markers along C) hair cell and D) supporting cell trajectories. *fat1a* and *pard3bb* only decrease along the hair cell trajectory, while *fgfr2* and *igsf3* decrease along both hair cell and supporting cell trajectories. E) Transient expression of early hair cell genes *dla* and *atoh1a* along hair cell trajectories. F) Increases in gene expression levels of *gfi1aa* and *myo6b* along hair cell trajectories. G) Increases in *stm* and *lfng* along supporting cell trajectories.

### Distinct supporting cell types in the cristae versus maculae

Supporting cells comprise two major clusters that can be distinguished by expression of *tectb* and *zpld1a* among other genes (Figure 3C, see Supplemental Table 4 for differentially expressed genes). The *tectb* gene encodes Tectorin beta, a component of the tectorial membrane associated with cochlear hair cells in mammals (Goodyear et al., 2017), and a component of otoliths in zebrafish (Kalka et al., 2019). The z*pld1a* gene, encoding Zona-pellucida-like domain containing protein 1a, is expressed in the cristae in fish (Dernedde et al., 2014; Yang et al., 2011) and mouse (Vijayakumar et al., 2019). Using fluorescent in situ hybridization, we find that *tectb* is expressed in the macular organs but is absent from cristae (Figure 5C). Conversely, *zpld1a* is expressed in cristae but not maculae (Figure 5D). In addition, *tectb* and *zpld1a* were not detected in lateral line neuromasts, showing they are inner ear-specific genes. Both *tectb* and *zpld1a* are expressed primarily in supporting cells, as they show little overlap in expression with the hair cell marker *myo6b*:GFP, similar to expression of the supporting cell marker *lfng* (Figure 5B-D, Supplemental Figure 4). These results demonstrate the presence of distinct supporting cell subtypes for the maculae and cristae.

**Figure 5.**
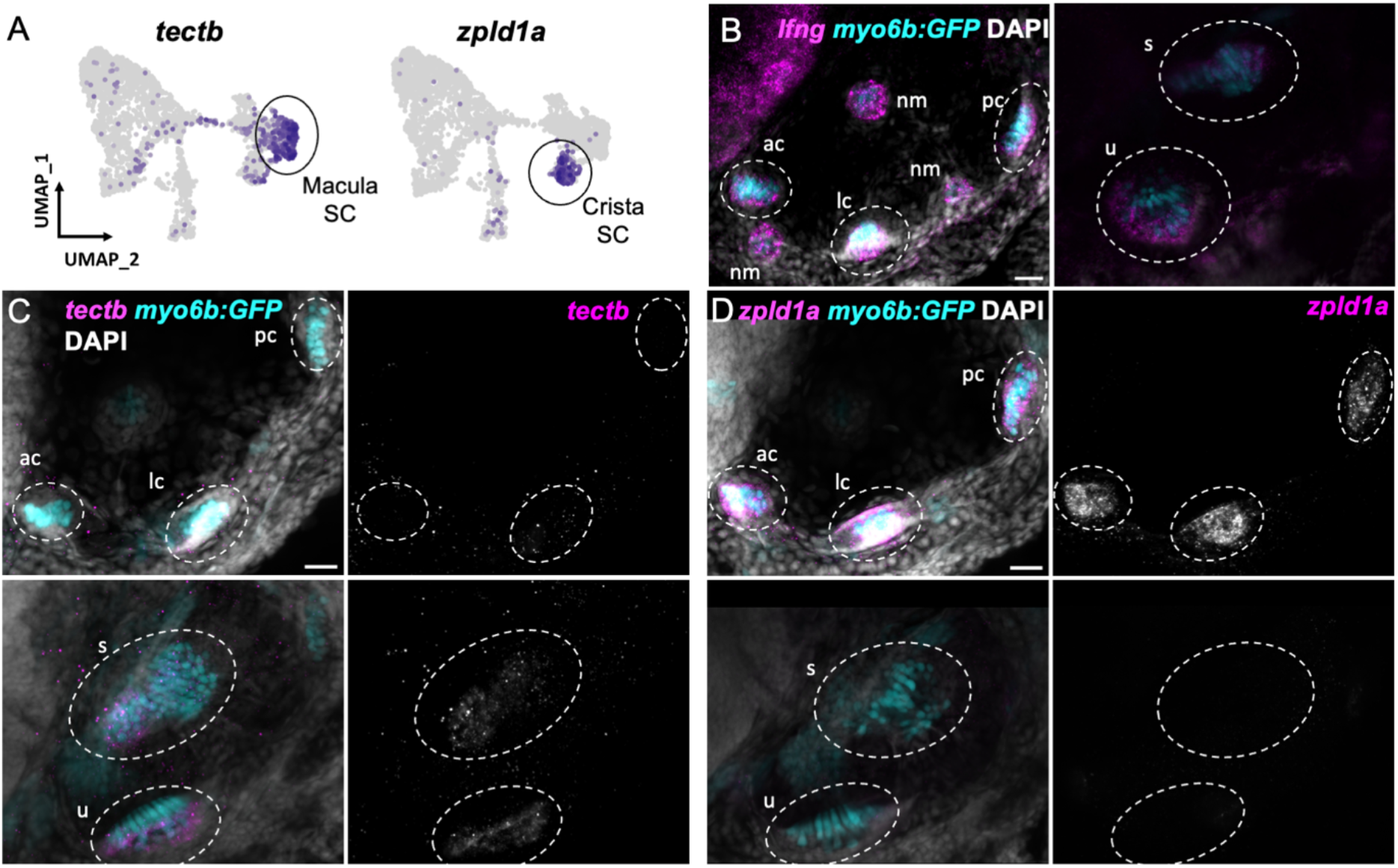
Distinct markers separate macula and crista supporting cells. A) Feature plots showing expression of macula supporting cell marker *tectb* and crista supporting cell marker *zpld1a*. B-D) HCR in situ hybridization in myo6b:GFP transgenic animals. Each set of images shown represents a projection of one z-stack split into cristae (lateral) and macula (medial) slices. Lateral line neuromasts positioned over the ear are visible in lateral slices. Expression pattern for B) the pan-supporting cell marker *lfng*, C) macula-specific marker *tectb*, and D) crista-specific marker *zpld1a* in 5 dpf myo6b:GFP fish. Each set of images shown represents a projection of one z-stack split into cristae (lateral) and macula (medial) slices. Lateral line neuromasts positioned over the ear are visible in lateral slices. ac: anterior crista, lc: lateral crista, nm: neuromast, pc: posterior crista, u: utricle, s: saccule. Scale bar = 20 μm.

### Distinct types of hair cells in the zebrafish inner ear

While inner ear and lateral line hair cells share many structural and functional features, we sought to determine if these cells also have distinct molecular signatures. We compared published datasets of lateral line hair cells (Baek et al., 2022; Kozak et al., 2020; Ohta et al., 2020) to our data, restricting analysis to datasets generated by 10x Chromium preparation to avoid technical batch effects across studies. Using Scanorama for alignments (Hie et al., 2019), hair cells from the inner ear and lateral line form distinct clusters, with a number of differentially expressed genes (Supplemental Figure 5), including the known markers for lateral line (*s100t*) and inner ear (*strc*) (Figure 3). This analysis suggests that inner ear hair cells of the maculae and cristae are more similar to each other than to lateral line hair cells.

Within the maculae and cristae, we find that hair cells can be subdivided into two major groups (clusters 1 and 3 versus cluster 4), distinguished by expression of two calcium binding protein genes, *cabp1b* and *cabp2b* (Di Donato et al., 2013) (Figure 3C). Hair cell cluster 5 has a mixed identity with co-expression of *cabp1b* and *cabp2b*. In the larval cristae, utricle, and saccule, *cabp1b* and *cabp2b* mark *myo6b*+ hair cells in largely non-overlapping zones (Figure 6B-D). By adult stages, complementary domains of *cabp1b*+ and *cabp2b*+ hair cells become clearly apparent. In the adult utricle, a central crescent of *cabp2b+; myo6b*+ hair cells is surrounded by a broad domain of *cabp1b*+; *myo6b*+ hair cells (Figure 6E, H). In the saccule and lagena, a late developing sensory organ, central *cabp2b+; myo6b*+ hair cells are surrounded by peripheral *cabp1b+; myo6b*+ hair cells (Figure 6F,G,I,J). We also find several genes that are specific for hair cells in the cristae, utricle, or saccule (Figure 7A). These include the calcium binding protein gene *cabp5b* in the cristae, the transcription factor *skor2* in the utricle, and the deafness gene *loxhd1b* in the saccule (Figure 7B-D, Supplemental Figure 6B).

**Figure 6.**
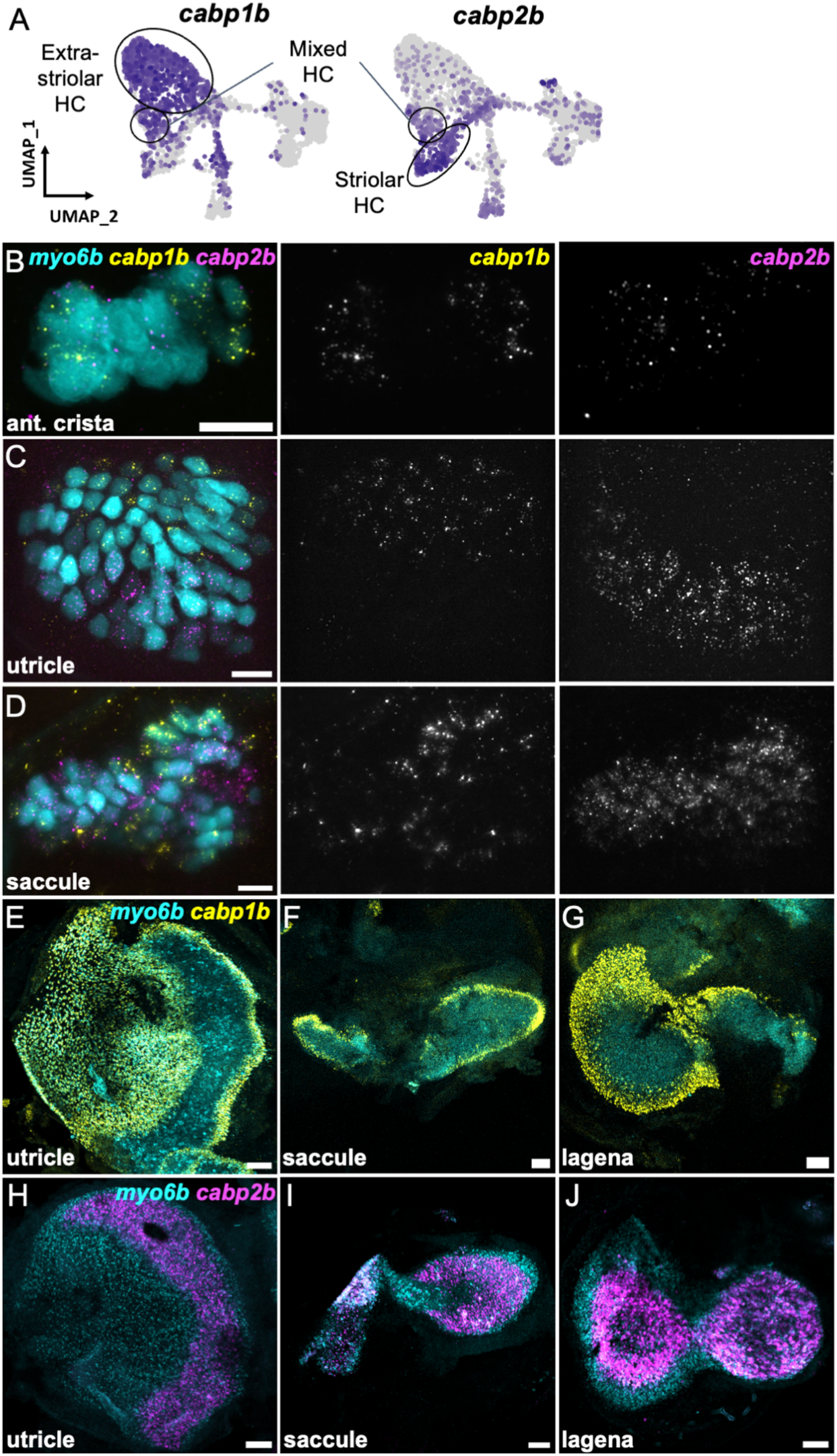
*cabp1b+* and *cabp2b*+ label hair cells in distinct regions of sensory end organs. A) Feature plots showing differential expression of *cabp1b* and *cabp2b* among crista and macula hair cells. B-D) HCR in situ projections of individual sensory patches from 5 dpf myo6:GFP fish showing differential spatial expression patterns of *cabp1b* and *cabp2b*. B) *cabp1b* is expressed at the ends of the cristae, while *cabp2b* is expressed centrally. Anterior crista is shown. C) In the utricle, *cabp1b* is expressed medially and *cabp2b* is expressed laterally. D) In the saccule, *cabp1b* is expressed in peripheral cells at the dorsal and ventral edges of the organ. *cabp2b* is expressed centrally. Scale bars for HCR images = 10 μm. E-G) Whole mount RNAScope confocal images of adult inner ear organs showing peripheral expression pattern of *cabp1b* (n = 3) in the adult zebrafish E) utricle, F) saccule, and G) lagena. H-J) Whole mount RNAScope confocal images showing central expression pattern of *cabp2b* (n = 4) in the adult zebrafish H) utricle, I) saccule, and J) lagena. Scale bars for RNAScope images = 25 μm.

**Figure 7.**
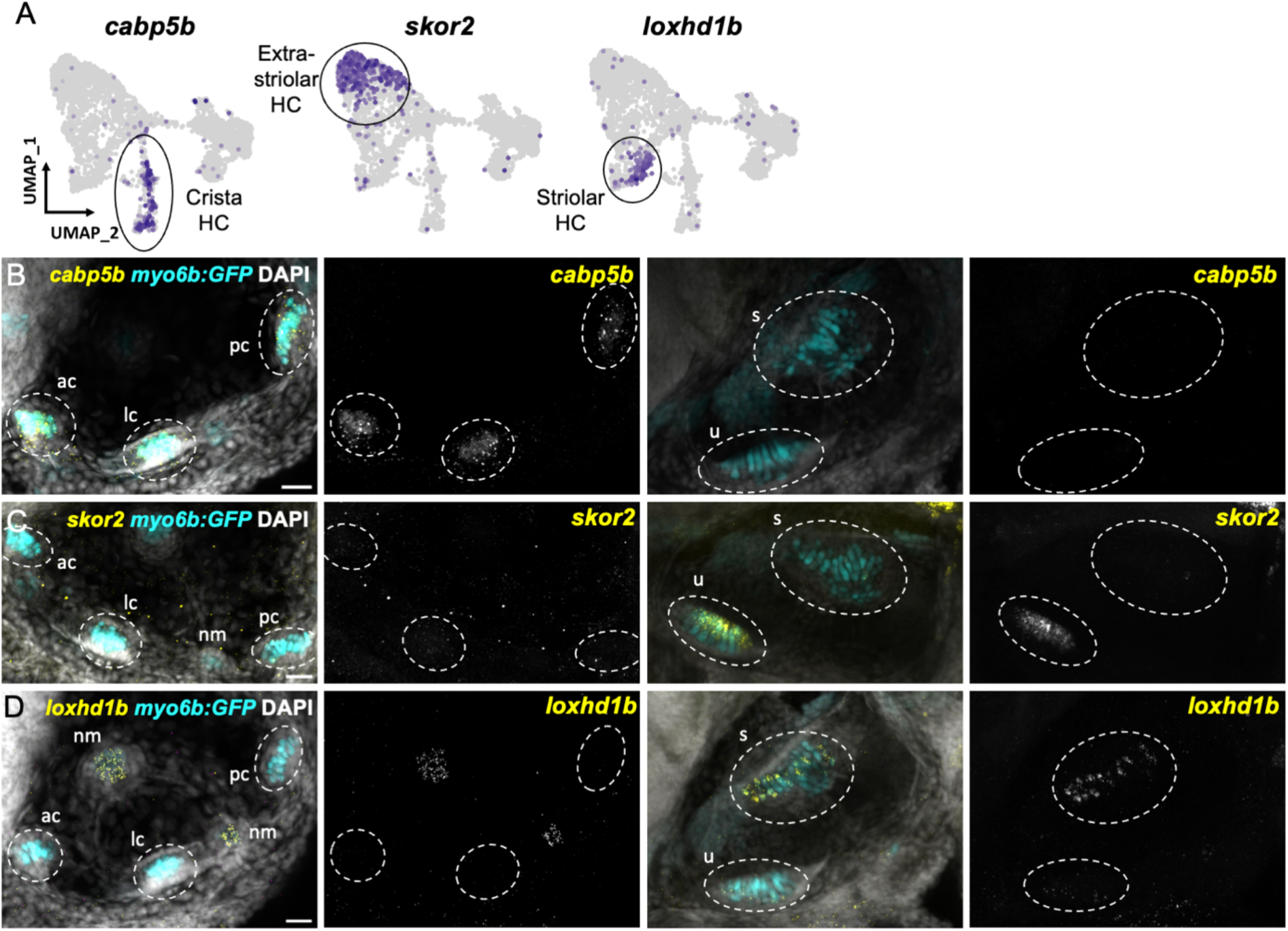
Distinct markers separate macula and crista hair cells. A) Feature plots showing marker genes enriched in organ-specific subsets of inner ear hair cells: *cabp5b, skor2*, and *loxhd1b*. B-D) HCR in situ for in 5dpf myo6b:GFP fish for genes shown in A). B) *cabp5b* is enriched in crista hair cells and is not expressed in macula hair cells. *skor2* is expressed in a subset of hair cells in the utricle only. D) *loxhd1b* is expressed in a subset of hair cells in the saccule. *loxhd1b* expression is also seen in hair cells of neuromasts near the ear. Each set of images shown represents an orthogonal projection of one z-stack split into cristae (lateral) and macular (medial) slices. ac: anterior crista, lc: lateral crista, nm: neuromast, pc: posterior crista, s: saccule, u: utricle. Scale bar = 20 μm.

The domain organization of *cabp2b*+ and *cabp1b+* hair cells in the adult macular organs resembles that of striolar and extrastriolar hair cells in the mammalian utricle. We therefore examined expression of *pvalb9*, the zebrafish ortholog of the mouse striolar hair cell marker *Ocm* (Hoffman et al., 2018; Jiang et al., 2017) (Figure 8, Supplemental Figure 7). In the larval utricle, we observe near complete overlap of *pvalb9* with *cabp2b* (Figure 8B). In the adult utricle, there is substantial overlap of *pvalb9* with *cabp2b* expression (except for a thin strip of *pvalb9*+; *cabp2b-* cells), and little overlap with *cabp1b* expression (Figure 8C,D). In addition, anti-Spectrin staining of hair bundles reveals a line of polarity reversal within the *cabp2b+* domain of the utricle (Figure 8E), consistent with polarity reversal occurring within the striolar domains of mammalian macular organs (Li et al., 2008). The *cabp2b*+ and *cabp1b*+ populations also differentially express genes related to stereocilia tip link and mechanotransduction channel components (Supplemental Figure 8, Supplemental Table 5) and various calcium and potassium channels (Supplemental Figure 9, Supplemental Table 5). These findings suggest that zebrafish *cabp2b*+ and *cabp1b*+ hair cells largely correspond to striolar and extrastriolar hair cells, respectively, with distinct mechanotransduction and synaptic properties.

**Figure 8.**
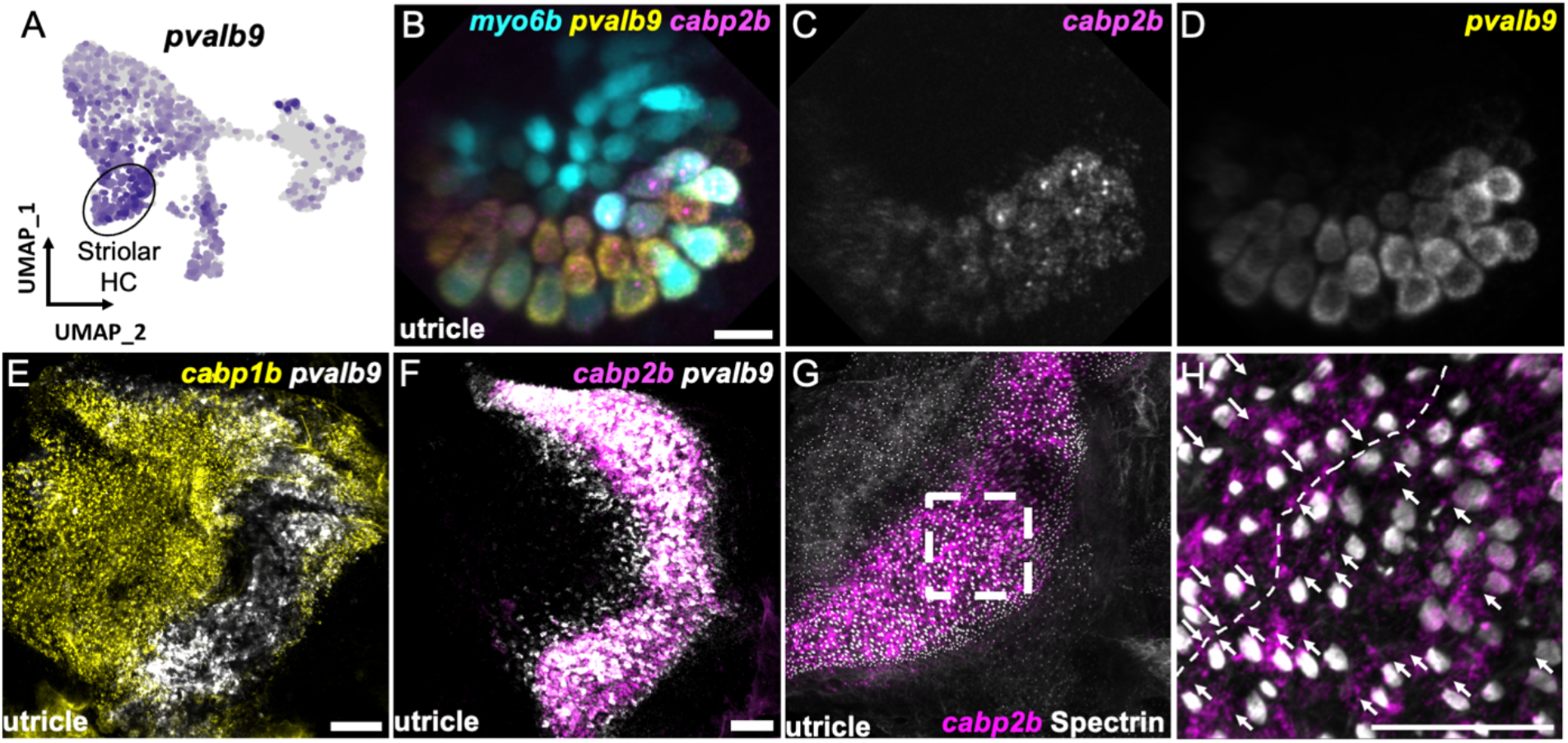
Zebrafish *cabp2b*+ domain coincides with features of mouse striolar region. A) Feature plot showing enrichment for the striola marker *pvalb9* in *cabp2b-expressing* cells. B-HCR in situ from 5dpf myo6b:GFP fish showing *pvalb9* and *cabp2b* expression in the utricle. Scale bar = 10 μm. E-F) Whole mount RNAScope confocal images of adult zebrafish utricles showing *pvalb9* probed alongside E) *cabp1b* (n = 3) and F) *cabp2b* (n = 4). Scale bar = 25 μm. G-H) Whole mount RNAScope RNA-protein co-detection assay showing co-localization of *cabp2b* expression (RNA) and hair cell line of polarity reversal indicated by Spectrin (protein) staining (n = 3). Scale bar = 25 μm. Arrows denote hair cell polarity and dotted line outlines line of polarity reversal.

**Figure 9.**
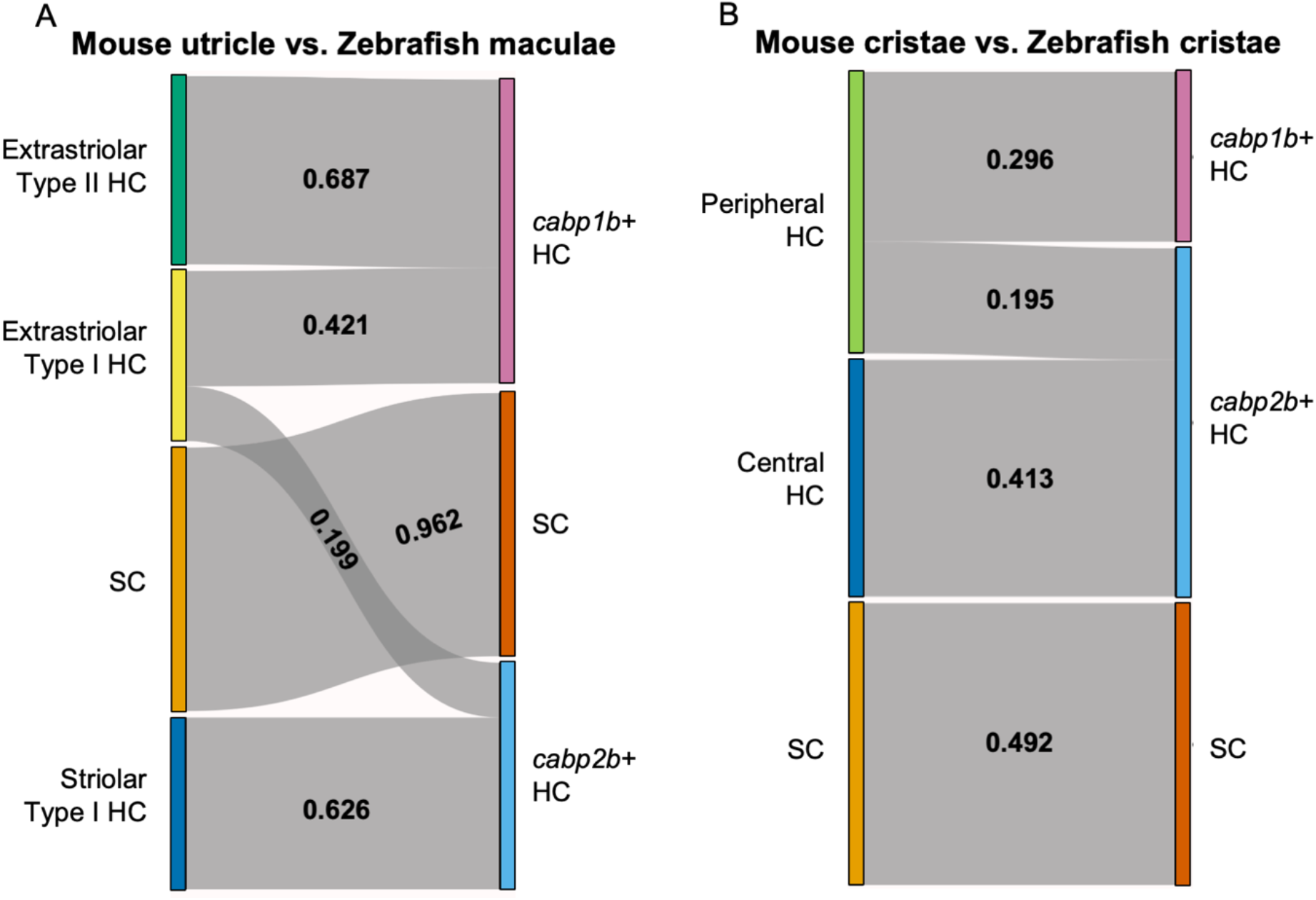
SAMap analysis reveals conserved gene expression patterns between mouse and zebrafish hair cell types. A-B) Sankey plot showing the SAMap mapping scores (0-1) that indicate transcriptome relatedness between A) mouse utriclar and zebrafish macular single-cell clusters and B) mouse and zebrafish cristae single-cell clusters. A mapping score of 0 indicates no evolutionary correlation in transcriptome while a mapping score of 1 indicates perfect correlation. Correlations below 0.15 were not plotted.

### Global homology of striolar and extrastriolar hair cells between fish and mice

To further probe similarities between zebrafish *cabp2b*+ and *cabp1b+* hair cells versus striolar and extrastriolar hair cells in mammals, we utilized the Self-Assembling Manifold mapping (SAMap) algorithm (Tarashansky et al., 2021; Musser et al., 2021) to compare cell types across distant species. A strength of this algorithm is that it compares not only homologous gene pairs but also close paralogs, which is especially useful considering the extensive paralog switching observed between vertebrate clades (Postlethwait, 2007), as well as the extra round of genome duplication in the teleost lineage leading to zebrafish. When comparing adult zebrafish maculae with the postnatal mouse utricle (Jan et al., 2021), we find the highest alignment score between supporting cells (Figure 9A). Consistent with the spatial domains revealed by our in situ gene expression analysis, we find that mouse striolar Type I hair cells exclusively map to zebrafish *cabp2b+* hair cells, and mouse extrastriolar Type I and Type II hair cells predominantly to zebrafish *cabp1b*+ hair cells. The small degree of mapping of mouse extrastriolar Type I hair cells to zebrafish *cabp2b+* hair cells suggests that zebrafish *cabp2b+* hair cells may have more of a Type I identity than *cabp1b+* cells in general. Gene pairs driving the alignment include striolar markers *Ocm, Loxhd1*, and *Atp2b2* for zebrafish *cabp2b+* hair cells, and mouse extrastriolar markers *Tmc1, Atoh1*, and *Jag2* for zebrafish *cabp1b+* hair cells (Supplemental Table 6).

A recent single-cell study revealed distinct central versus peripheral hair cell subpopulations in postnatal mouse cristae, reminiscent of the striolar and extrastriolar populations in the maculae (Wilkerson et al., 2021). As our zebrafish cristae hair cells also separate by the expression of *cabp1b* and *cabp2b* (Figure 6A), we performed SAMap analysis between the crista cell populations of the two species to investigate cell type homology. Similar to what we observed for the utricle, zebrafish centrally located *cabp2b+* crista hair cells predominantly map to mouse central crista hair cells, and zebrafish peripherally located *cabp1b+* crista hair cells exclusively map to mouse peripheral crista hair cells (Figure 9B, see Supplemental Tables 7 and 8 for differentially expressed genes in *cabp1b*+ and *cabp2b*+ crista hair cells and gene pairs that are driving homology). In contrast, zebrafish lateral line hair cells (Lush et al., 2019) align exclusively to mouse extrastriolar and not striolar hair cells (Supplemental Figure 10). Thus, zebrafish *cabp2b+* macular hair cells are closely related to striolar cells of the mouse utricle, with zebrafish lateral line and *cabp1b+* macular hair cells more closely related to mouse extrastriolar hair cells.

## Discussion

Our single-cell transcriptomic profiling of the embryonic to adult zebrafish inner ear reveals a diversity of hair cell and supporting cell subtypes that differ from those of the lateral line. We identify hair cells and supporting cells specific for each inner ear end organ, as well as two spatially segregated types of zebrafish inner ear hair cells with similarities to mammalian striolar and extrastriolar hair cells. The identification of end organ-specific hair cells and supporting cell types points to different functions of each macular organ that will be interesting to pursue in the future. Moreover, as much of our knowledge about zebrafish hair cell regeneration comes from studies of the lateral line, understanding similarities and differences between the lateral line and inner ear has the potential to uncover mechanisms underlying the distinct regenerative capacity of inner ear hair cell subtypes. Recent tools to systematically damage inner ear hair cells in zebrafish (Jimenez et al., 2021) should enable such types of comparative studies.

Our study shows that zebrafish possess distinct types of striolar and extrastriolar hair cells in the maculae and cristae, with molecular differences between these subtypes implying different physiological properties. In each of the end organs, the two subtypes of hair cells are defined by differential expression of calcium binding proteins, in particular *cabp1b* versus *cabp2b*. As calcium binding proteins closely interact with synaptic ion channels (Pitcher et al., 2017), their differential expression may confer unique electrophysiological properties. In support of this idea, these hair cell subtypes express distinct combinations of ion channel genes and mechanotransduction components (Supplemental Figs 8 and 9). Our findings are consistent with previous reports of distinct current profiles in central versus peripheral hair cells in the zebrafish utricle, saccule, and lagena (Haden et al., 2013; Olt et al., 2014), as well as spatial differences in ciliary bundle morphology and synaptic innervation in the larval zebrafish utricle (Liu et al., 2022). The distinct spatial distribution, channel expression, and hair bundle morphologies in these hair cells resembles the known spatial, electrophysiological, and hair bundle compositional differences seen in the striolar versus extrastriolar hair cells in the amniote vestibular end organs (Holt et al., 2007; Kharkovets et al., 2000; Lapeyre et al., 1992; Meredith and Rennie, 2016; Moravec and Peterson, 2004; Rüsch et al., 1998; Xue and Peterson, 2006). Our work extends this to show extensive molecular similarities between the central *cabp2b+* and peripheral *cabp1b+* hair cells and mammalian striolar and extrastriolar hair cells, respectively.

By contrast, we found no clear homology of zebrafish inner ear hair cells with mammalian Type I and Type II hair cells. The lack of molecular signatures corresponding to Type I hair cells is consistent with previous reports that one of their major features, calyx synapses, are absent from fishes (Lysakowski and Goldberg, 2004). These findings suggest that the diversification of inner ear hair cells into Type I and Type II cells emerged after the evolutionary split of ray-finned fishes from the lineage leading to mammals.

Our integrated dataset reveals distinct molecular characteristics of hair cells and supporting cells in the zebrafish inner ear sensory organs, with conservation of these patterns from larval stages to adults. In addition, although not discussed in detail here, our data include additional cell populations of the zebrafish inner ear that express extracellular matrix-associated genes important for otic capsule structure and ion channel-associated genes associated with fluid regulation. These data form a resource that can be further explored to inform molecular aspects of hair cell electrophysiology, mechanotransduction, sound versus motion detection, maintenance of inner ear structure and ionic balance, and inner ear-specific hair cell regeneration.

## Materials and Methods

### Zebrafish lines

This study was performed in strict accordance with the recommendations in the Guide for the Care and Use of Laboratory Animals of the National Institutes of Health. The Institutional Animal Care and Use Committees of the University of Southern California (Protocol 20771) and University of Washington (Protocol 2997-01) approved all animal experiments. Experiments were performed on zebrafish (*Danio rerio*) of AB or mixed AB/Tubingen background. For adult stages, mixed sexes of animals were used for constructing single-cell libraries, as well as RNAScope experiments. Published lines include *Tg(Mmu*.*Sox10-Mmu*.*Fos:Cre)*^*zf384*^ (Kague et al., 2012); *Tg(−3*.*5ubb:LOXP-EGFP-LOXP-mCherry)*^*cz1701Tg*^ (Mosimann et al., 2011); and *Tg(myosin 6b:GFP)*^*w186*^ (Hailey et al., 2017).

### In situ hybridization and RNAScope

Hybridization chain reaction in situ hybridizations (Molecular Instruments, HCR v3.0) were performed on 5 dpf *myo6b*:GFP larvae as directed for whole-mount zebrafish embryos and larvae (Choi et al., 2016, 2018). Briefly, embryos were treated with 1-phenyl 2-thiourea (PTU) beginning at 24 hpf. At 5 dpf, larvae were fixed in 4% PFA overnight at 4°C. Larvae were washed with PBS and then stored in MeOH at -20°C until use. Larvae were rehydrated using a gradation of MeOH and PBST washes, treated with proteinase K for 25 min and post-fixed with 4% PFA for 20 min at room temperature. For the detection phase, larvae were pre-hybridized with a probe hybridization buffer for 30 min at 37°C, then incubated with probes overnight at 37°C. Larvae were washed with 5X SSCT to remove excess probes. For the amplification stage, larvae were pre-incubated with an amplification buffer for 30 min at rt and incubated with hairpins overnight in the dark at rt. Excess hair pins were removed by washing with 5X SSCT. Larvae were treated with DAPI and stored at 4°C until imaging. All HCR in situ patterns were confirmed in at least 3 independent animals. Transcript sequences submitted to Molecular Instruments for probe generation are listed in Supplemental Table 9. The *cabp1b* probes were tested on 3 separate occasions and imaged in at least 6 animals; *cabp2b* probes were tested on 5 separate occasions and imaged in at least 20 different animals; *cabp5b* probes were tested on 3 separate occasions and imaged in at least 9 different animals; *lfng* probes were tested on two separate occasions and imaged in at least 5 different animals; *loxhd1b* probes were tested on two separate occasions and imaged in at least 7 animals; *pvalb9* probes were tested on two separate occasions and imaged in at least 6 different animals; *skor2* probes were tested on two separate occasions and imaged in at least 6 different animals; *tectb* probes were tested on 4 separate occasions and imaged in at least 10 different animals; *zpld1a* probes were tested on 3 separate occasions and imaged in at least 9 different animals.

RNAScope samples were prepared by fixation in 4% paraformaldehyde either at room temperature for 2 hours or at 4 °C overnight. Adult (28-33mm) inner ears were dissected and dehydrated in methanol for storage. RNAScope probes were synthesized by Advanced Cell Diagnostics (ACD): Channel 1 probe *myo6b* (1045111-C1), Channel 2 probe *pvalb9* (1174621-C2), and Channel 3 probes *cabp1b* (1137731-C3) and *cabp2b* (1137741-C3). Whole inner ear tissues were processed through the RNAScope Fluorescent Multiplex V2 Assay (ACD Cat. No. 323100) according to manufacturer’s protocols with the ACD HybEZ Hybridization oven. *cabp1b* probe was tested on 4 separate occasions with 6 animals or 12 ears total; *cabp2b* probe was tested on 4 separate occasions with 7 animals or 14 ears total; *pvalb9* probe was tested on 2 separate occasions with 6 animals or 12 ears total. *myo6b* probe was used with each of the above probes.

### Immunofluorescence staining

Immediately following the RNAScope protocol, samples were prepared for immunofluorescence staining using mouse anti-β-Spectrin II antibody (BD Bioscience Cat. No. 612562, RRID: AB_399853). Briefly, RNAScope probed zebrafish ears were rehydrated in PBS for 5 min and rinsed in PBDTx (0.5 g bovine serum albumin, 500 μL DMSO, 250 μL 20% Triton-X in 50 mL PBS, pH = 7.4) for 15 min at room temperature. They were then blocked in 2% normal goat serum (NGS) in PBDTx for 3 hours at room temperature, and incubated with 1:500 dilution of mouse anti-β-Spectrin II antibody in PBDTx containing 2% NGS overnight at 4 °C. After 3 washes in PBDTx for 20 min each at room temperature, samples were incubated with 1:1000 dilution of Alexa 647 goat-anti-mouse IgG1 secondary antibody (Invitrogen Cat. No. A-21240, RRID: AB_2535809) for 5 hours at room temperature. They were then washed 2 times in PBSTx (250 μL 20% Triton-X in 50 mL PBS) for 5 min each before imaging. Three animals or 6 ears total were subjected to Spectrin detection on 2 separate occasions.

### Imaging

Confocal images of whole-mount RNAScope samples were captured on a Zeiss LSM800 microscope (Zeiss, Oberkochen, Germany) using ZEN software. HCR-FISH imaging was performed on a Zeiss LSM880 microscope (Zeiss, Oberkochen, Germany) with Airyscan capability. Whole larvae were mounted between coverslips sealed with high vacuum silicone grease (Dow Corning) to prevent evaporation. Z-stacks were taken through the ear at intervals of 1.23 μm using a 10X objective or through individual inner ear organs at an interval of 0.32 μm using a 20X objective. 3D Airyscan processing was performed at standard strength settings using Zen Blue software.

### Single-cell preparation and analysis

#### scRNA-seq library preparation and alignment

For 14 dpf animals (n=35), heads from converted *Sox10:Cre; ubb:LOXP-EGFP-LOXP-mCherry* fish were decapitated at the level of the pectoral fin with eyes and brains removed. For 12 mpf animals (n=6, 27-31mm), utricle, saccule, and lagena were extracted from converted *Sox10:Cre; ubb:LOXP-EGFP-LOXP-mCherry* fish after brains and otolith crystals were removed. Dissected heads and otic sensory patches were then incubated in fresh Ringer’s solution for 5–10 min, followed by mechanical and enzymatic dissociation by pipetting every 5 min in protease solution (0.25% trypsin (Life Technologies, 15090-046), 1 mM EDTA, and 400 mg/mL Collagenase D (Sigma, 11088882001) in PBS) and incubated at 28.5 °C for 20–30 min or until full dissociation. Reaction was stopped by adding 6× stop solution (6 mM CaCl2 and 30% fetal bovine serum (FBS) in PBS). Cells were pelleted (376 × g, 5 min, 4 °C) and resuspended in suspension media (1% FBS, 0.8 mM CaCl2, 50 U/mL penicillin, and 0.05 mg/mL streptomycin (Sigma-Aldrich, St. Louis, MO) in phenol red-free Leibovitz’s L15 medium (Life Technologies)) twice. Final volumes of 500 μL resuspended cells were placed on ice and fluorescence-activated cell sorted (FACS) to isolate live cells that excluded the nuclear stain DAPI. For scRNAseq library construction, barcoded single-cell cDNA libraries were synthesized using 10X Genomics Chromium Single Cell 3′ Library and Gel Bead Kit v.3.1 (14 dpf) or Single Cell Multiome ATAC + Gene Expression kit (12 mpf, ATAC data not shown) per the manufacturer’s instructions. Libraries were sequenced on Illumina NextSeq or HiSeq machines at a depth of at least 1,000,000 reads per cell for each library. Read2 was extended from 98 cycles, per the manufacturer’s instructions, to 126 cycles for higher coverage. Cellranger v6.0.0 (10X Genomics) was used for alignment against GRCz11 (built with GRCz11.fa and GRCz11.104.gtf) and gene-by-cell count matrices were generated with default parameters.

#### Data processing of scRNA-seq

Count matrices of inner ear and lateral line cells from embryonic and larval timepoints (18-96 hpf) were analyzed using the R package Monocle3 (v1.0.0) (Cao et al., 2019). Matrices were processed using the standard Monocle3 workflow (preprocess_cds, detect_genes, estimate_size_factors, reduce_dimension(umap.min_dist = 0.2, umap.n_neighbors = 25L)). This cell data set was converted to a Seurat object for integration with 10X Chromium sequencing data using SeuratWrappers. The count matrices of scRNA-seq data (14 dpf and 12 mpf) were analyzed by R package Seurat (v4.1.0) (Hao et al., 2021). Cells of neural crest origins were removed bioinformatically based on our previous study (Fabian et al., 2022). The matrices were normalized (NormalizeData) and integrated with normalized scRNA-seq data from the embryonic and larval time points according to package instruction (FindVariableFeatures, SelectIntegrationFeatures, FindIntegrationAnchors, IntegrateData; features = 3000). The integrated matrices were then scaled (ScaleData) and dimensionally reduced to 30 principal components. The data were then subjected to neighbor finding (FindNeighbors, k = 20) and clustering (FindClusters, resolution = 0.5), and then visualized through UMAP with 30 principal components as input. After data integration and processing, RNA raw counts from all matrices were normalized and scaled according to package instructions to determine gene expression for all sequenced genes, as the integrated dataset only contained selected features for data integration.

Mouse utricle scRNA-seq data (Jan et al., 2021) was downloaded from NCBI Gene Expression Omnibus (GSE155966). The count matrix was analyzed by R package Seurat (v4.1.0). Matrices were normalized (NormalizeData) and scaled for the top 2000 variable genes (FindVariableFeatures and ScaleData). The scaled matrices were dimensionally reduced to 15 principal components. The data were then subjected to neighbor finding (FindNeighbors, k = 20) and clustering (FindClusters, resolution = 1) and visualized through UMAP with 15 principal components as input. Hair cells and supporting cells were bioinformatically selected based on expression of hair cells and supporting cell markers *Myo6* and *Lfng*, respectively. Hair cells were further subcategorized into striola type I hair cells by co-expression of striola marker *Ocm* and type I marker *Spp*, extrastriola type I hair cells by expression of *Spp* without *Ocm*, and extrastriola type II hair cells by expression of *Anxa4* without *Ocm*.

Mouse crista scRNA-seq data (Wilkerson et al., 2021) was downloaded from NCBI Gene Expression Omnibus (GSE168901). The count matrix was analyzed by R package Seurat (v4.1.0). Matrices were normalized (NormalizeData) and scaled for the top 2000 variable genes (FindVariableFeatures and ScaleData). The scaled matrices were dimensionally reduced to 15 principal components. The data were then subjected to neighbor finding (FindNeighbors, k = 20) and clustering (FindClusters, resolution = 1) and visualized through UMAP with 15 principal components as input. Hair cells and supporting cells were bioinformatically selected based on expression of hair cell and supporting cell markers *Pou4f3* and *Sparcl1*, respectively. Hair cells were further subcategorized into central hair cells by expression of *Ocm* and peripheral hair cells by expression of *Anxa4*.

#### Pseudotime analysis

We used the R package Monocle3 (v1.0.1) to predict the pseudo temporal relationships within the integrated scRNA-seq dataset of sensory patches from 36 hpf to 12 mpf. Cell paths were predicted by the learn_graph function of Monocle3. We set the origin of the cell paths based on the enriched distribution of 36 to 48 hpf cells. Hair (all macular hair cells, clusters 0-5) and supporting (macular supporting cells clusters 0 and 6) cell paths were selected separately (choose_cells) to plot hair cells and supporting cell marker expression along pseudotime (plot_genes_in_pseudotime).

#### Differential gene expression

We utilized *presto* package’s differential gene expression function to identify differentially expressed genes among the different cell types. Wilcox rank sum test was performed by the function *wilcox usc*. We then filtered for genes with log2 fold change greater than 0.5 and adjusted p-value less than 0.01. To compare inner ear hair cells to lateral line hair cells, we used the following datasets from GEO: GSE144827 (Kozak et al., 2020), GSE152859 (Ohta et al., 2020), and GSE196211(Baek et al., 2022). Hair cells were selected from datasets by expression of *otofb* and integrated along with our 10x Chromium dataset with Scanorama (Hie et al., 2019).

#### SAMap analysis for cell type homology

We used the python package SAMap (v1.0.2)(Tarashansky et al., 2021) to correlate gene expression patterns and determine cell type homology between mouse utricle (GSE155966) (Jan et al., 2021) or crista (GSE168901) (Wilkerson et al., 2021) hair cells and supporting cells and our 12 mpf zebrafish inner ear scRNA-seq data. Zebrafish lateral line hair cell sc-RNA data (GSE123241) (Lush et al., 2019) was integrated with our 12 mpf inner ear data using Seurat in order to compare to mice. First, a reciprocal BLAST result of the mouse and zebrafish proteomes was obtained by performing blastp (protein-protein BLAST, NCBI) in both directions using in-frame translated peptide sequences of zebrafish and mouse transcriptome, available from ensembl (Danio_rerio.GRCz11.pep.all.fa and Mus_musculus.GRCm38.pep.all.fa). The generated maps were then used for the SAMap algorithm. Raw count matrices of zebrafish and mouse scRNA-seq Seurat objects with annotated cell types were converted to h5ad format using SeuratDisk package (v0.0.0.9020) and loaded into Python 3.8.3. Raw data were then processed and integrated by SAMap. Mapping scores between cell types of different species were then calculated by get_mapping_scores and visualized by sankey_plot. Gene pairs driving cell type homology were identified by GenePairFinder.

## Data availability

Single cell RNA seq data are available from the NCBI Gene Expression Omnibus with Gene Set Accession number GSE211728.

## Acknowledgments

This manuscript is dedicated to Neil Segil, who was a wonderful colleague, friend, and mentor to many. We thank Megan Matsutani for fish care, the USC Stem Cell Flow Cytometry Core, and the CHLA Next-Generation Sequencing Core. We also thank David White and the UW Zebrafish Facility staff for fish care.

## Author Contributions

TS and MOB conceived, designed and performed experiments, performed data analysis, wrote the manuscript. PF performed experiments. LMS and CT provided data and guided analysis. NS, JGC and DWR conceived and designed experiments, performed data analysis, wrote the manuscript.

## Figures & Figure Legends

**Figure S1.**
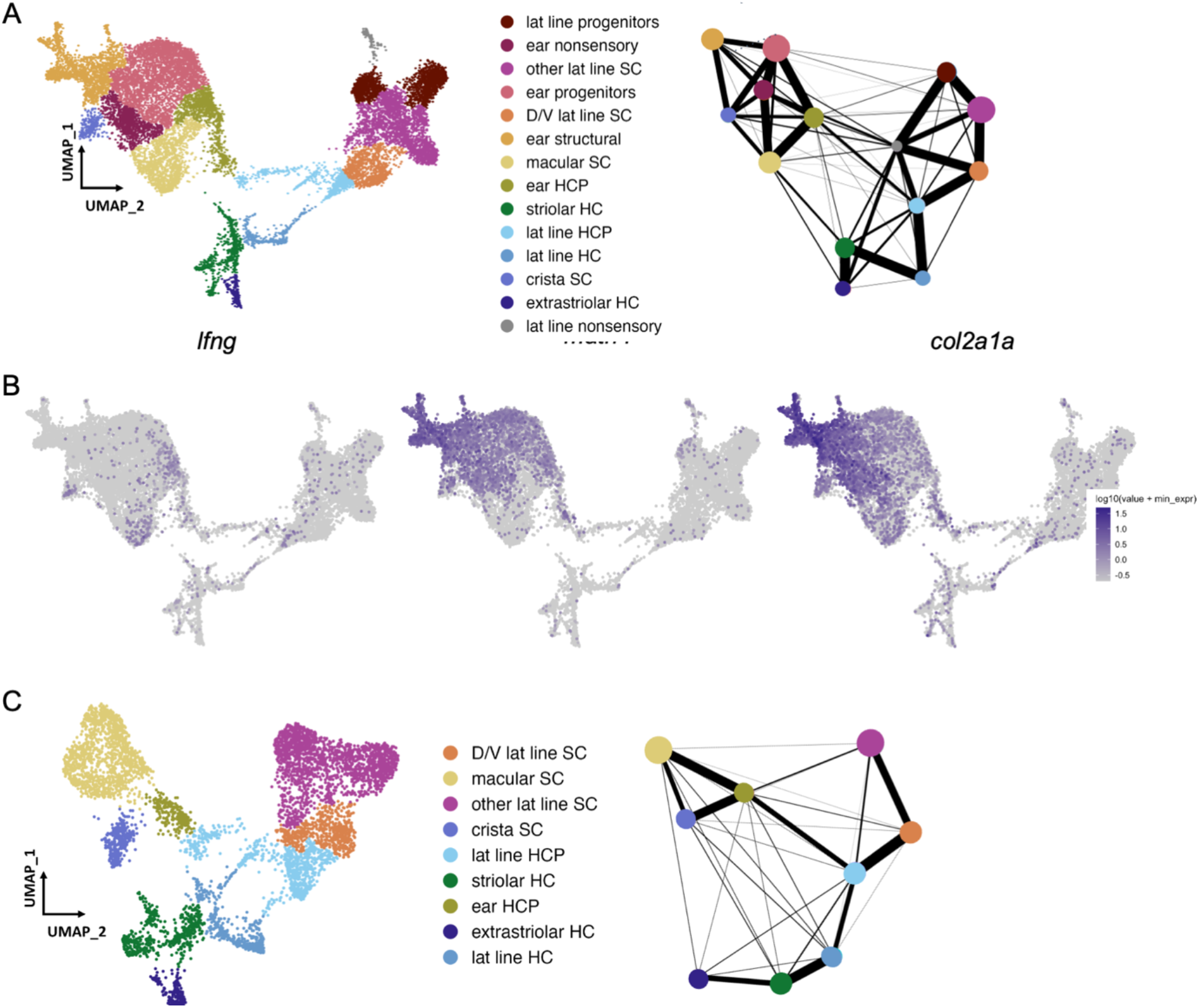
Selection of otic sensory cells from snRNA-seq dataset. A) Clustering of 18 hpf to 96 hpf dataset to illustrate cell subtypes. PAGA analysis of this dataset shows strong connectivity among ear nonsensory cells and lateral line nonsensory cells, but weak interconnectivity between these two groups. B) Feature plots showing expression of the supporting cell marker *lfng*, and markers of structural otic cells *matn4* and *col2a1a*. C) UMAP of sensory patch cells from 36-96hpf are-clustered without structural and early otic vesicle cells. PAGA analysis again shows strong connectivity within hair cells and supporting cell groups and weak connectivity between lateral line and inner ear supporting cells. PAGA connectivity scores are listed in Supplemental Table 1.

**Figure S2.**
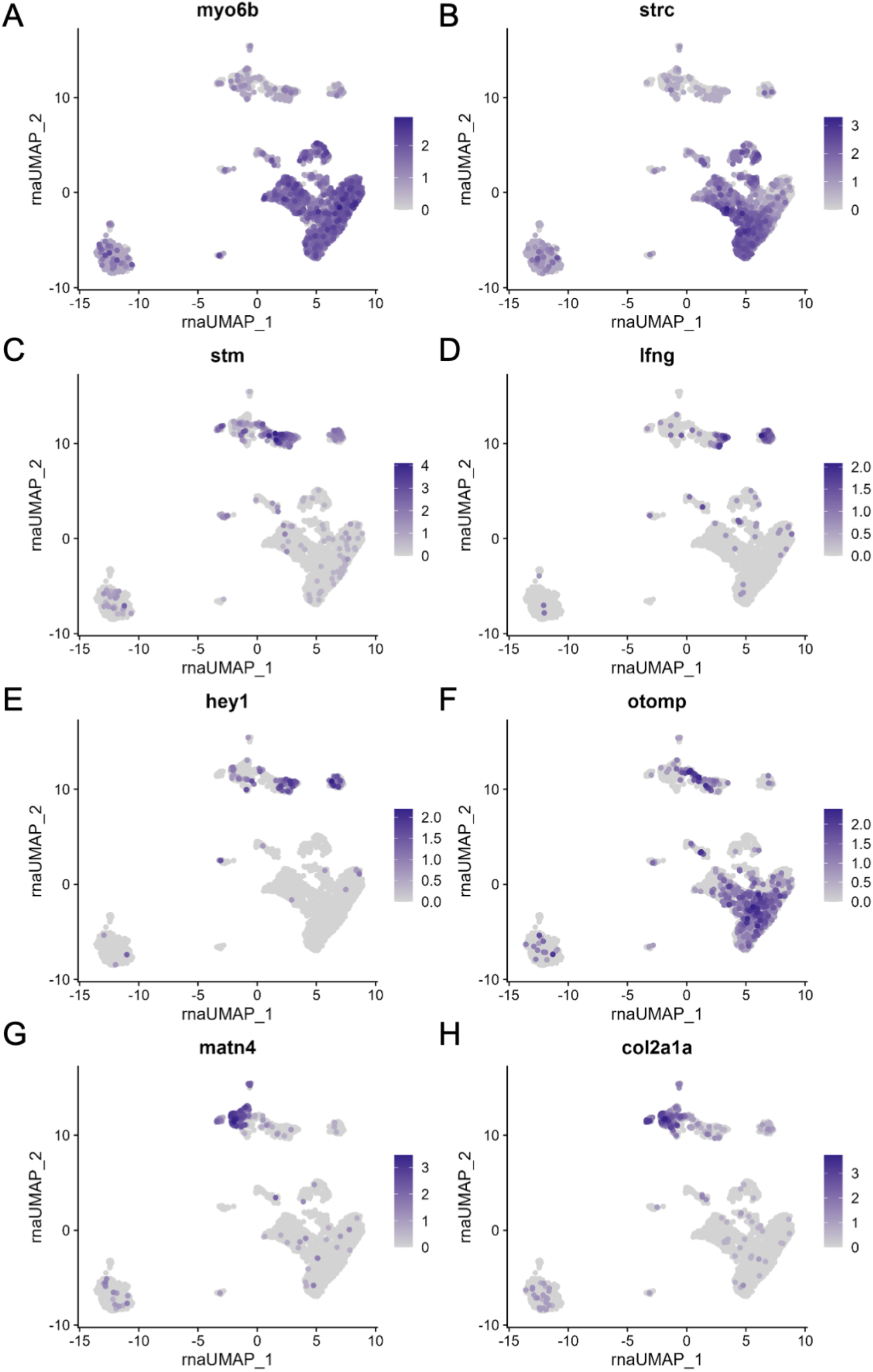
scRNA-seq of 12 mpf zebrafish inner ear captures sensory hair cells and supporting cells as well as non-sensory supporting cells. Feature plots of 12 mpf zebrafish scRNA-seq dataset alone showing expression of hair cell markers A) *myo6b* and B) *strc*, pan-supporting cell marker C) *stm*, sensory supporting cell markers D) *lfng* and E) *hey1*, and pan-otic marker F) *otomp*, and non-sensory supporting cell markers G) *matn4*, and H) *col2a1a*.

**Figure S3.**
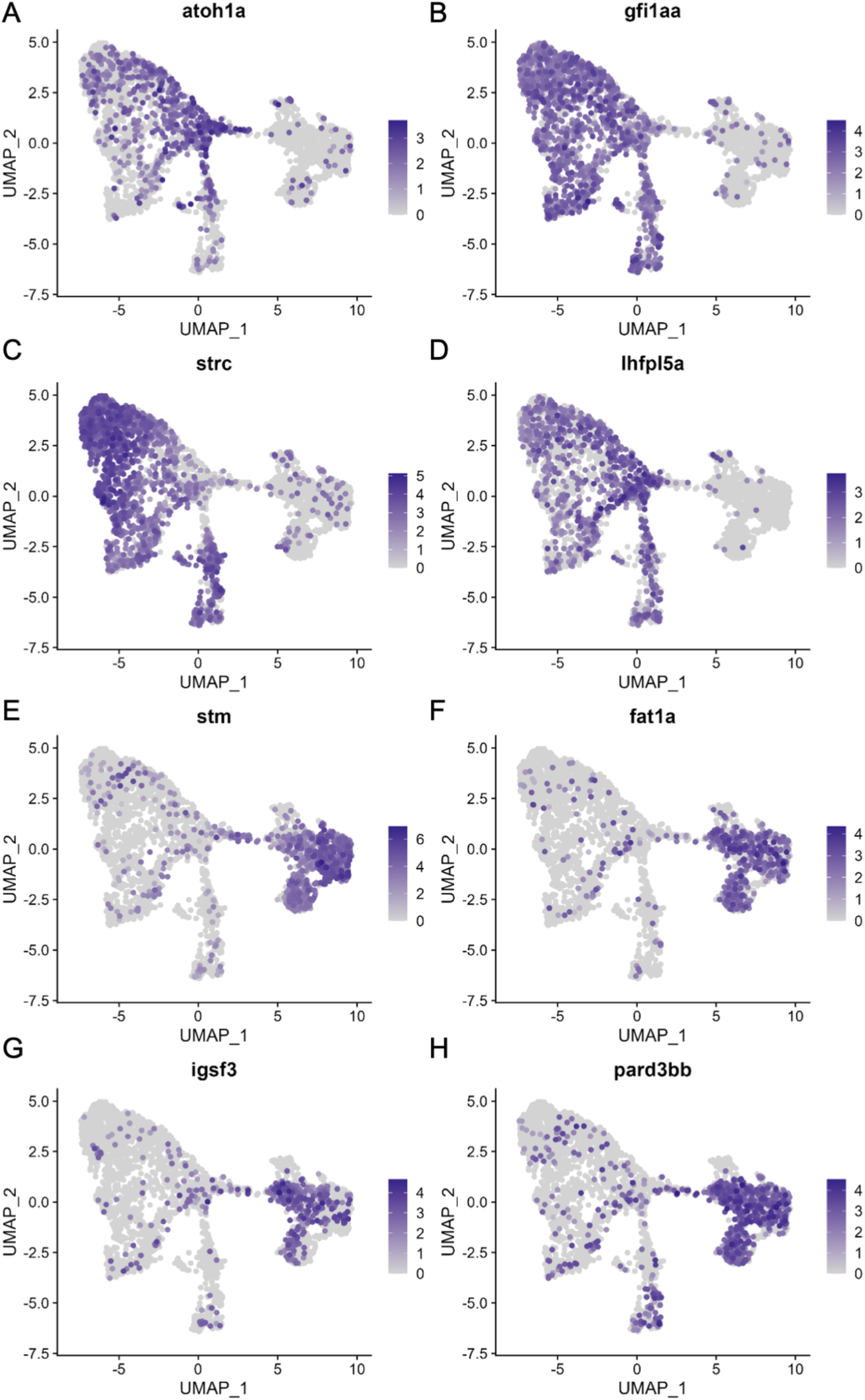
hair cell and supporting cell marker expression in the integrated scRNA-seq dataset. Feature plots of integrated zebrafish scRNA-seq datasets showing expression of nascent hair cell marker A) *atoh1a*, inner hair cell markers b) *strc*, c) *gfi1aa* and D) *lhfpl5a*, pan-supporting cell marker E) *stm*, and putative progenitor markers F) *fat1a*, G) *igsf3* and H) *pard3bb*.

**Figure S4.**
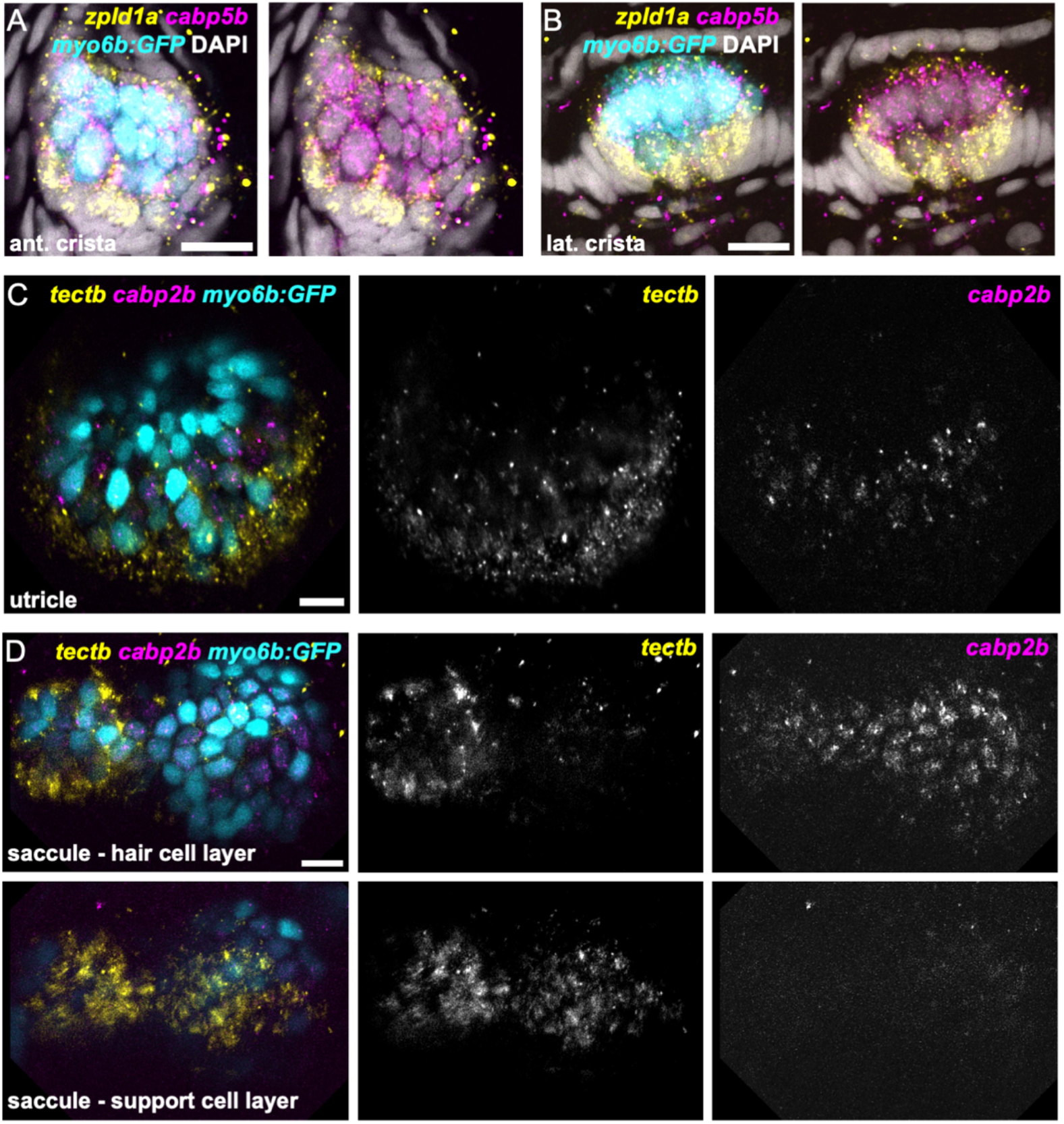
*zpld1a* and *tectb* are primarily expressed in supporting cells. HCR in situ hybridization of 5 dpf myo6b:GFP zebrafish. A-B) Confocal slice through A) anterior crista and B) lateral crista (lateral view) showing localization of *cabp5b* and *zpld1a* expression. *cabp5b* is primarily expressed in hair cells and zpld1a is primarily expressed in supporting cells. C) Slice through utricle (dorsal view) showing *cabp2b* expression in hair cells and *tectb* expression primarily in the surrounding supporting cells. D) Slices through saccule (lateral view) at the level of hair cell bodies (top row) and supporting cell bodies (bottom row). *cabp2b* is primarily expressed in hair cells and *tectb* is primarily expressed in supporting cells. Scale bars = 10 μm.

**Figure S5.**
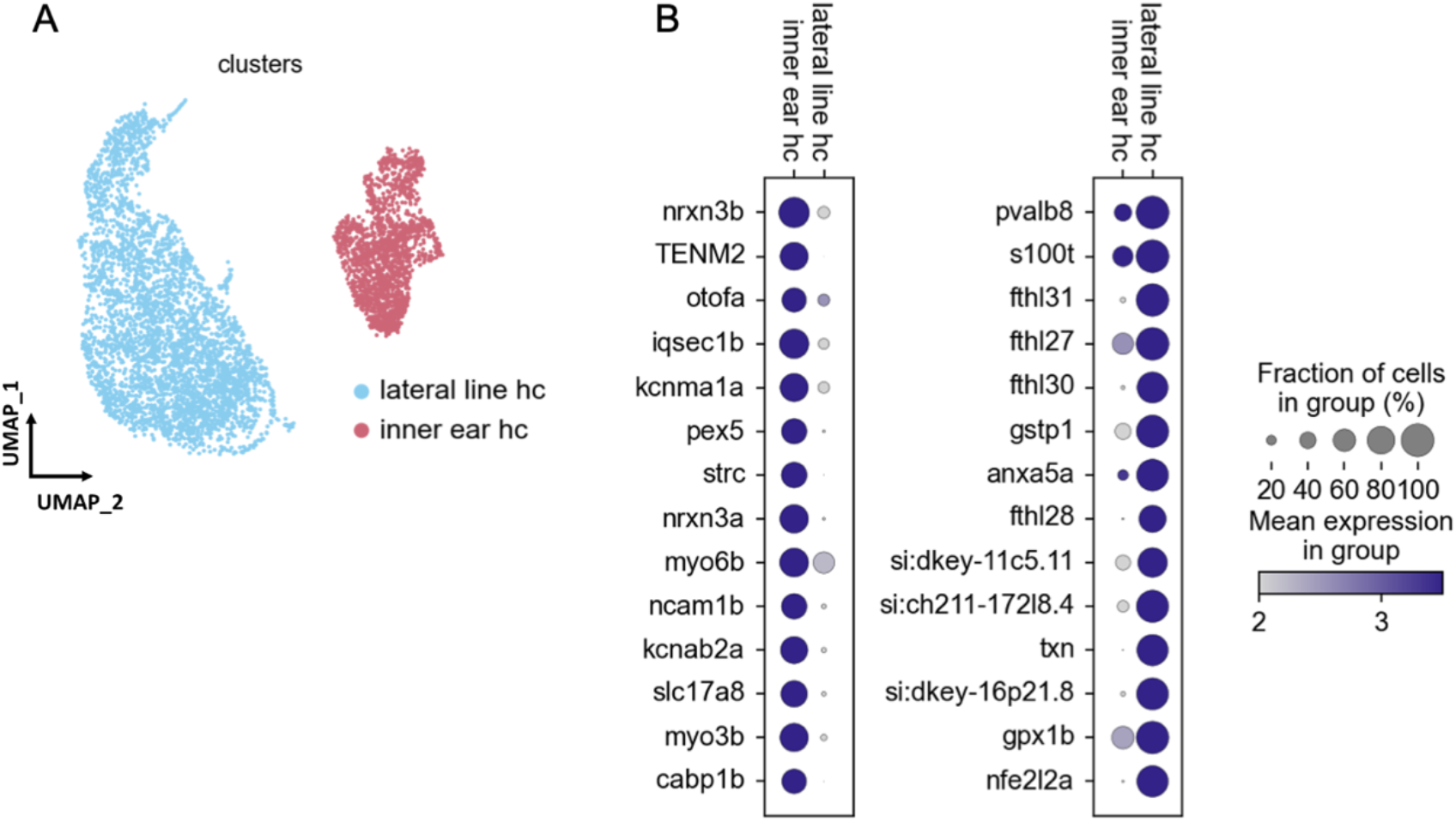
Gene expression differences between lateral line and inner ear hair cells. A) UMAP of hair-cell dataset comprising 12mpf cells from our dataset as well as lateral line hair cells from previously published datasets integrated with Scanorama. Lateral line hair cells cluster separately from inner ear hair cells. B) Differential gene expression analysis identifies novel marker genes specific to either lateral line or inner ear hair cells.

**Figure S6.**
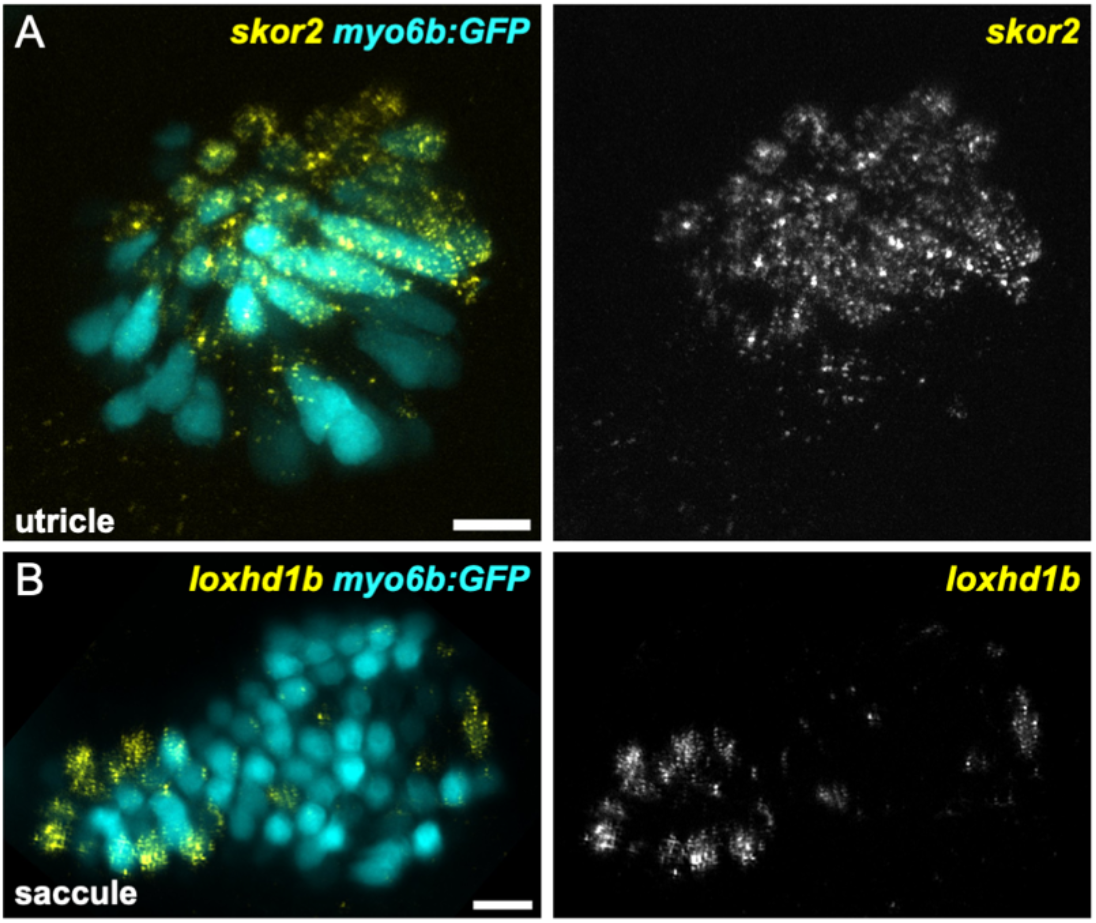
*skor2* and *loxhd1b* label subsets of hair cells in utricle or saccule. HCR in situ hybridization of 5 dpf zebrafish. A) Maximum intensity projection of utricle (dorsal view) showing *skor2* expression in medially-located hair cells. B) Maximum intensity projection of saccule (lateral view) showing *loxhd1b* expression in a peripheral subset of hair cells. Scale bars = 10 μm.

**Figure S7.**
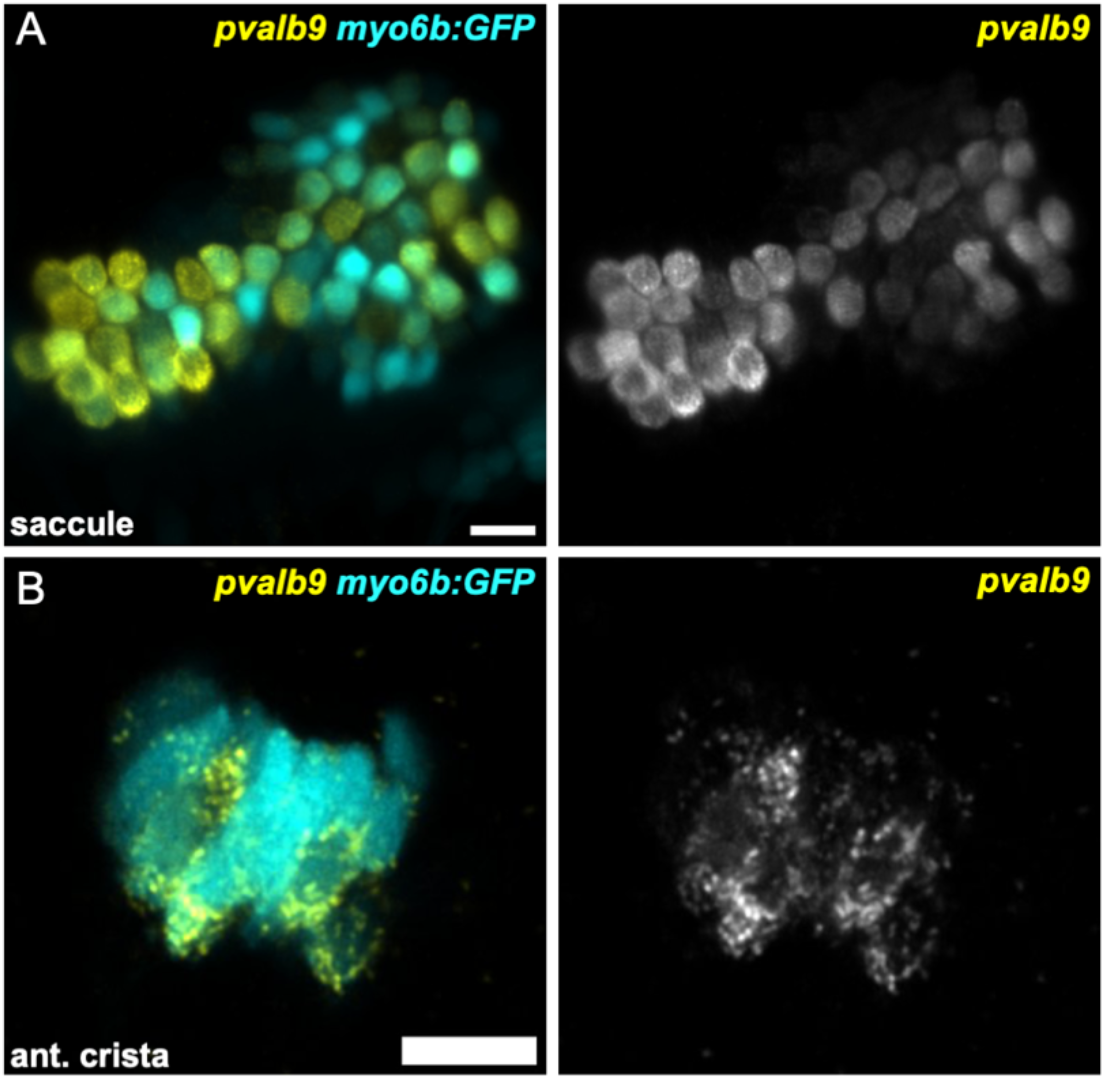
Striola marker *pvalb9* is expressed in all inner ear sensory end organs. HCR in situ hybridization of 5 dpf zebrafish. A) Maximum intensity projection of saccule (lateral view) showing *pvalb9* expression in centrally-located hair cells. B) Slice through the anterior crista showing *pvalb9* expression in a subset of crista hair cells. Scale bars = 10 μm.

**Figure S8.**
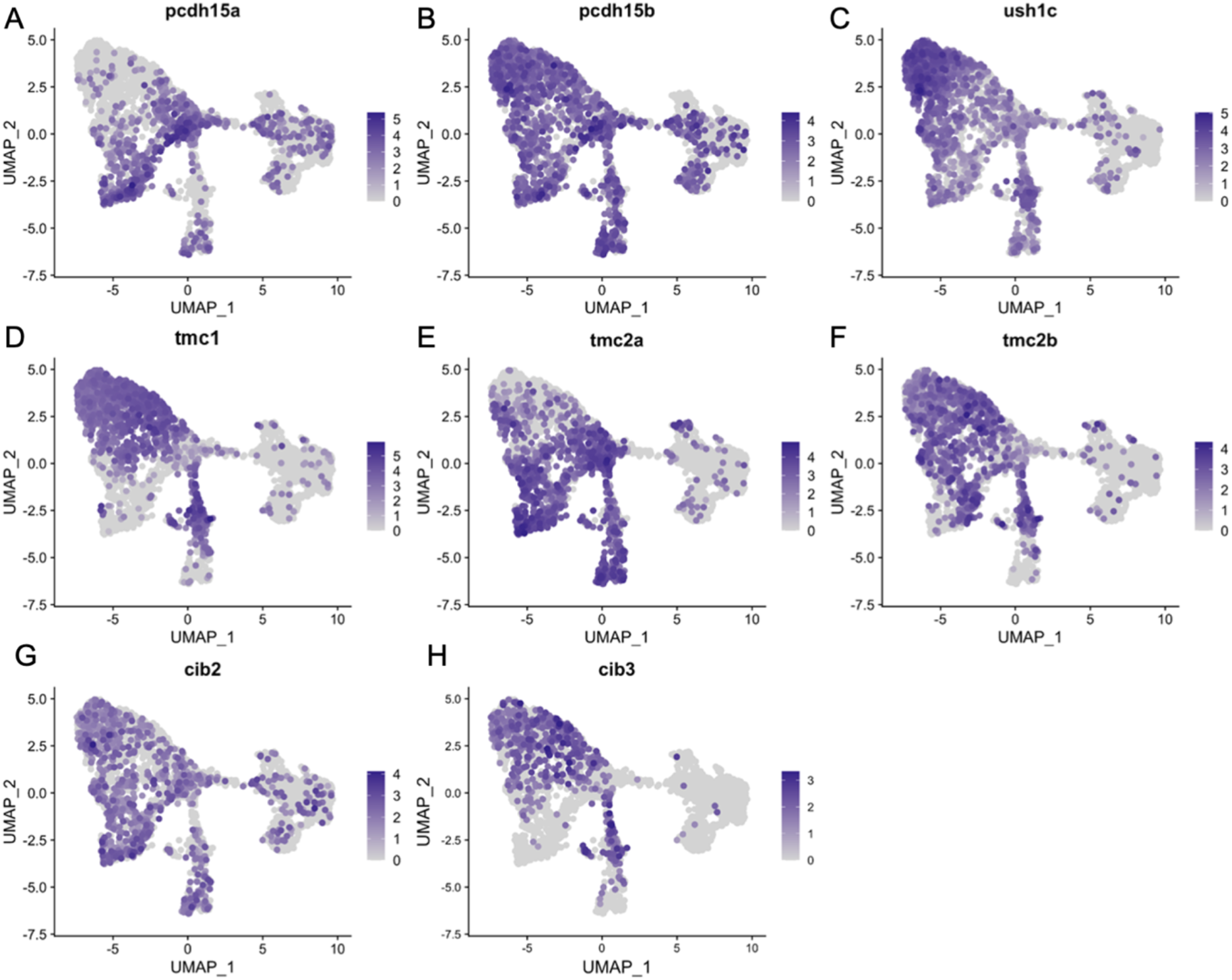
*cabp1b*+ and *cabp2b*+ hair cells differentially express mechanosensory apparatus genes. Feature plots of integrated zebrafish scRNA-seq datasets showing expression of A) *pcdh15a*, B) *pcdh15b*, C) *ush1c*, D) *tmc1*, E) *tmc2a*, F) *tmc2b*, G) *cib2*, and H) *cib3*.

**Figure S9.**
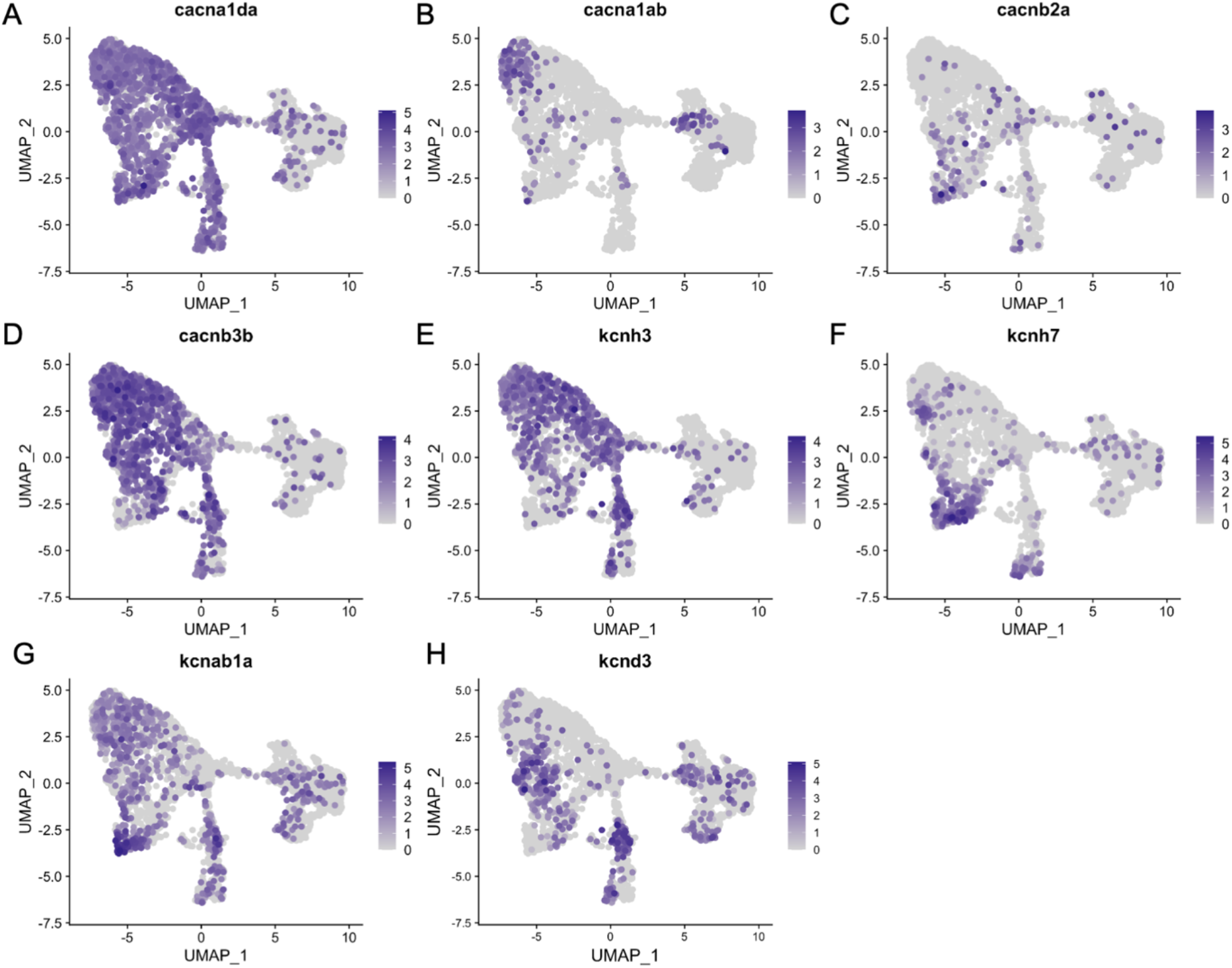
*cabp1b*+ and *cabp2b*+ hair cells differentially express voltage-gated calcium and potassium channel genes. Feature plots of integrated zebrafish scRNA-seq datasets showing expression of voltage-gated calcium channel genes A) *canada*, B) *cacna1ab*, C) *cacnb2a*, D) *cacnb3b*, and voltage-gated potassium channel genes E) *kcnh3*, F) *kcnh7*, G) *kcnaba*, and H) *kcnd3*.

**Figure S10.**
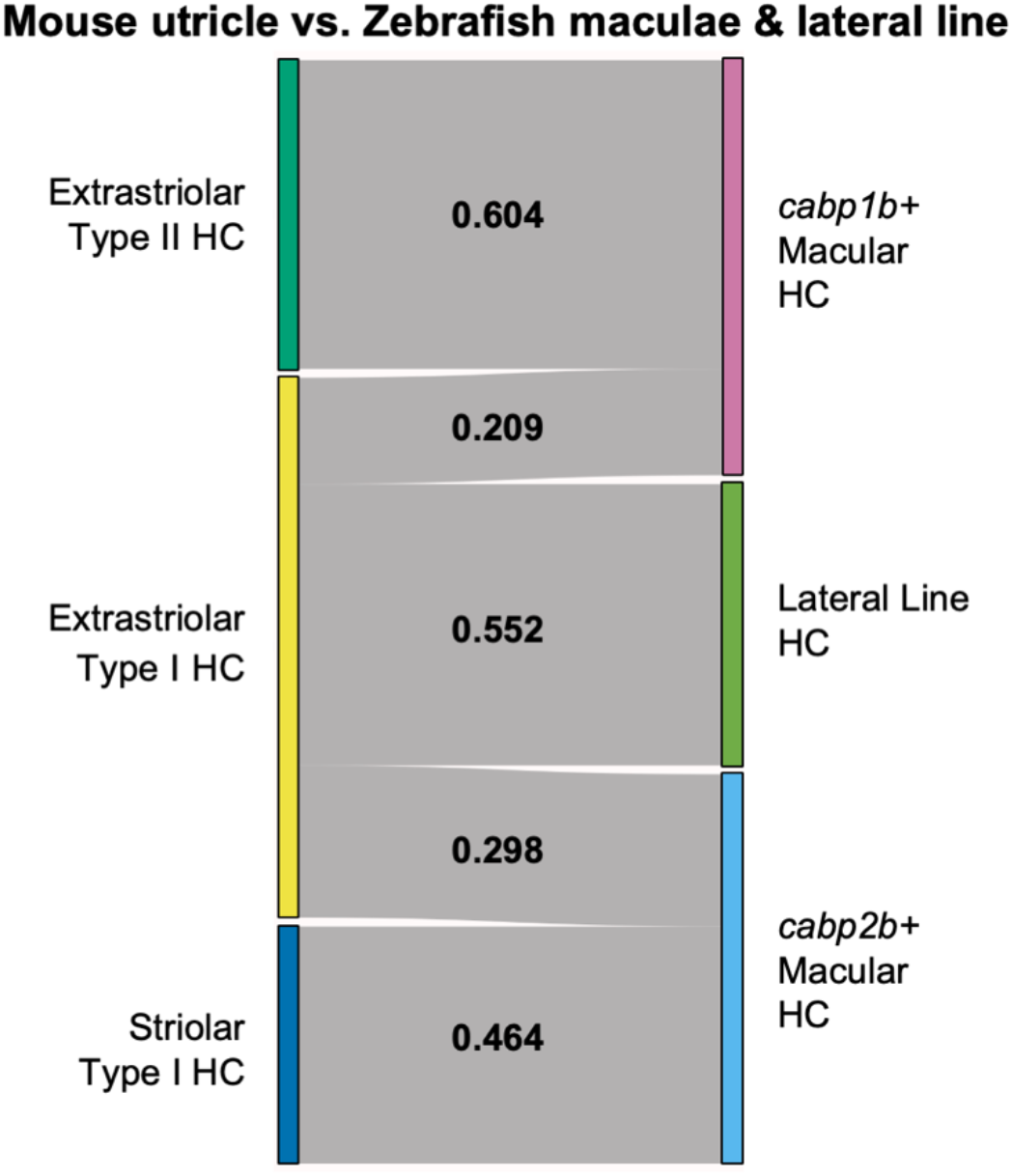
SAMap analysis of mouse utricle versus zebrafish macular and lateral line hair cells and supporting cells. A) Sankey plot showing the SAMap mapping scores (0-1) that indicate transcriptome relatedness between mouse utricular and integrated zebrafish macular and lateral line single-cell clusters. A mapping score of 0 indicates no evolutionary correlation in transcriptome while a mapping score of 1 indicates perfect correlation. Correlations below 0.2 were not plotted.

## List of Tables

Supplemental Table 1. PAGA scores for relative connectivity between clusters (related to Supplemental Figure 1)

Supplemental Table 2. Differentially expressed genes in inner ear cell clusters

Supplemental Table 3. Genes enriched along pseudotime trajectories

Supplemental Table 4. Genes enriched in supporting cell clusters

Supplemental Table 5. Genes enriched in *cabp1b+* and *cabp2b+* macular cells

Supplemental Table 6. Genes driving macular SAMap alignment

Supplemental Table 7. Genes enriched in *cabp1b+* and *cabp2b+* crista cells

Supplemental Table 8. Genes driving crista SAMap alignment

Supplemental Table 9. cDNA sequences used for HCR in situ hybridization probes

